# Accurate *de novo* detection of somatic mutations in high-throughput single-cell profiling data sets

**DOI:** 10.1101/2022.11.22.517567

**Authors:** Francesc Muyas, Ruoyan Li, Raheleh Rahbari, Thomas J. Mitchell, Sahand Hormoz, Isidro Cortés-Ciriano

## Abstract

Characterization of somatic mutations at single-cell resolution is essential to study cancer evolution, clonal mosaicism, and cell plasticity. However, detection of mutations in single cells remains technically challenging. Here, we describe SComatic, an algorithm designed for the detection of somatic mutations in single-cell transcriptomic and ATAC-seq data sets without requiring matched bulk or single-cell DNA sequencing data. Using >1.5M single cells from 383 single-cell RNAseq and single-cell ATAC-seq data sets spanning cancer and non-neoplastic samples, we show that SComatic detects mutations in single cells, even in differentiated cells from polyclonal tissues not amenable to mutation detection using existing methods. In addition, SComatic permits the estimation of mutational burdens and *de novo* mutational signature analysis at single-cell and cell-type resolution. Notably, using matched exome and single-cell RNAseq data, we show that SComatic achieves a 20 to 40-fold increase in precision as compared to existing algorithms for somatic SNV calling without compromising sensitivity. Overall, SComatic opens the possibility to study somatic mutagenesis at unprecedented scale and resolution using high-throughput single-cell profiling data sets.

## Main

Characterization of somatic mutations at single-cell resolution is essential to study genetic heterogeneity and cell plasticity in cancer^1^, clonal mosaicism in non-neoplastic tissues^2^, and to identify the mutational processes operative in both malignant and phenotypically normal cells^3,4^. Single-cell genome sequencing provides the most direct way to study mutations in single cells. However, single-cell genomics methods are not easily scalable, and suffer from high rates of genomic drop-outs and artefacts introduced during whole-genome amplification^5^. To circumvent the issues associated with whole-genome amplification, other approaches rely on bulk sequencing of single-cell-derived colonies grown *in vitro* or clonal populations directly isolated from tissues^6–8^. However, *in vitro* growth of single-cell-derived colonies is laborious and limited to cell types amenable to cell culture^5,7,9^, and isolation of clonal units is not technically feasible for some tissues. More recently, the development of ultra-sensitive sequencing methods using strand-specific barcoding has permitted detection of mutations at single-molecule resolution, even in polyclonal tissues^10,11^. Yet, cell type information is lost unless cell sorting is performed prior to sequencing. Due to these technical limitations, our understanding of the patterns of somatic mutations across cell types and their impact on cell fates and phenotypes remains limited.

An alternative strategy consists of detecting somatic mutations in sequencing reads from high-throughput single-cell profiling assays directly, such as single-cell RNA-seq (scRNA-seq) and single-cell assay for transposase-accessible chromatin using sequencing (scATAC-seq). The main advantage of this approach is the possibility to harness the high throughput of single-cell profiling assays to map the lineage of cells to transcriptional or regulatory programmes^12,13^ without the need for complex experimental protocols for joint profiling of the DNA and RNA from the same cell^3,8,14–16^. Nevertheless, detection of mutations is strongly limited due to the variability in gene expression across cell types, allelic drop-out events, transcriptional bursts, RNA editing, limited depth of coverage, and sequencing artefacts^17–19^. Therefore, existing algorithms rely on detecting mutations, such as single-nucleotide variants (SNVs) or indels, previously identified using matched bulk or single-cell DNA sequencing data^18,20–22^. These approaches are limited because matched DNA sequencing data are rarely available for existing high-throughput single-cell data sets, and due to sampling biases or genetic heterogeneity between the samples undergoing DNA sequencing and single-cell profiling. Therefore, algorithms designed to detect somatic mutations in single-cell data sets *de novo* without requiring matched DNA sequencing data are critically needed.

To address this need, we developed SComatic, an algorithm for *de novo* detection of somatic SNVs in single-cell profiling data sets, including scRNA-seq and scATAC-seq data, without requiring matched bulk or single-cell DNA sequencing data. Using a total of 1,575,862 non-neoplastic and cancer cells from 317 scRNA-seq and 66 scATAC-seq published data sets (Supplementary Table 1), we show that SComatic achieves a 20 to 40-fold increase in precision as compared to existing algorithms for somatic SNV calling without compromising sensitivity. In addition, we show that SComatic permits the detection of mutational burdens and *de novo* discovery of mutational signatures at cell-type resolution, even for differentiated cells and cells from polyclonal tissues showing high levels of genetic heterogeneity, which are not amenable to mutation detection using existing experimental or computational methods. SComatic is implemented in Python 3 and is available at https://github.com/cortes-ciriano-lab/SComatic.

## Results

### Overview of SComatic

We developed SComatic to detect somatic mutations using single-cell sequencing data without requiring matched bulk or single-cell DNA sequencing data (Fig. 1). In brief, SComatic computes base counts for every position of the genome across cell types from the same individual using cell type annotations established through e.g., marker gene expression (Fig. 1 and Methods). Somatic mutations are distinguished from germline polymorphisms and artefacts using a set of hard filters and statistical tests (Fig. 1). Specifically, SComatic only considers genomic positions with coverage in at least 5 cells from at least 2 cell types. Candidate somatic SNVs are distinguished from background sequencing errors and artefacts using a Beta-binomial test parameterized using non-neoplastic samples (Methods). Next, mutations detected in multiple cell types are considered to be germline polymorphisms or artefacts and are thus discounted as somatic. The key idea is that germline variants should be present in all cell types, whereas somatic mutations should only be detected in cell types from the same differentiation hierarchy, unless mutations were acquired in a progenitor or stem cell prior to clonal diversification or during early development^8,23,24^. Candidate mutations overlapping known RNA editing sites or single-nucleotide polymorphisms (SNPs) with population frequencies greater than 1 % in gnomAD^25^ are also filtered out. In addition, SComatic uses a ‘Panel of Normals’ generated using a large collection of non-neoplastic samples to discount recurrent sequencing or mapping artefacts. For example, in 10x Chromium scRNA-seq data, recurrent errors are enriched in LINE and SINE elements, such as Alu elements (Supplementary Fig. 1), which are thus not considered for mutation calling. Finally, to make a mutation call, SComatic requires a sequencing depth of at least 5 reads in the cell type in which the mutation is detected, and that the mutation is detected in at least 3 sequencing reads from at least 2 different cells of the same type (Supplementary Fig. 2 and Methods).

**Figure 1.**
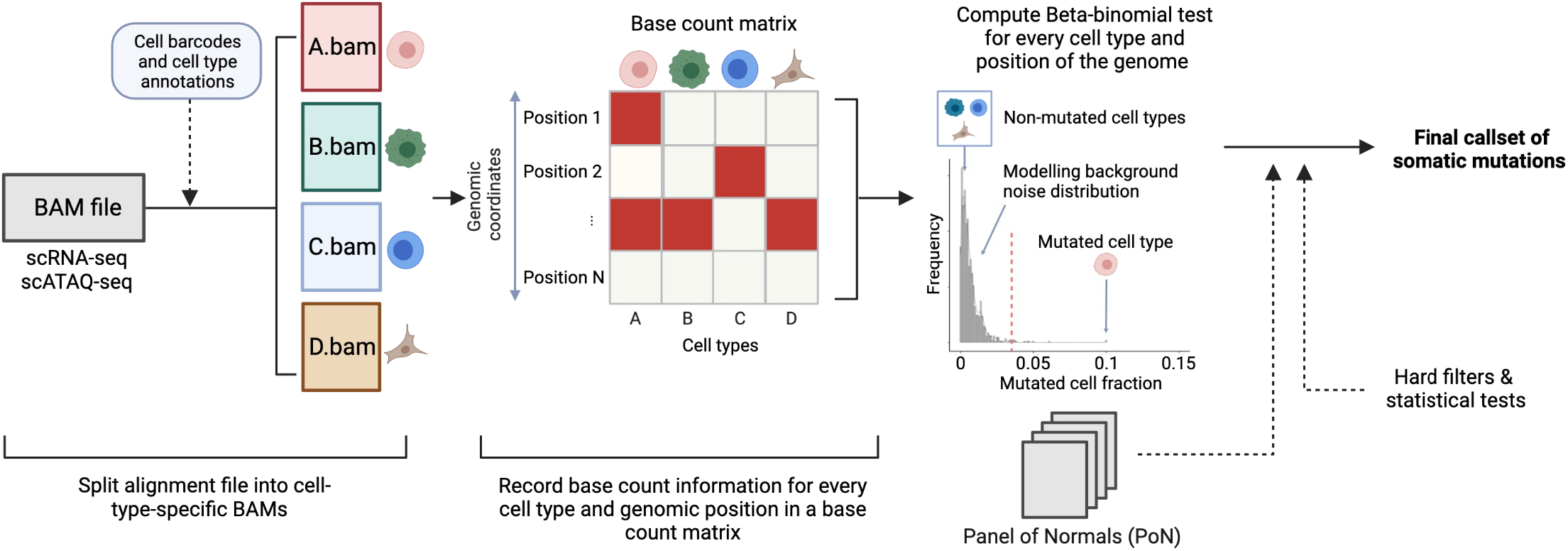
Overview of SComatic. Methodology for detecting somatic mutations in high-throughput single-cell profiling data sets.

### Validation of SComatic using matched single-cell RNA-seq and exome sequencing data

To compare the patterns of mutations detected by SComatic against DNA sequencing data, we analysed scRNA-seq data generated using the 10X Genomics Chromium technology and matched whole-exome sequencing (WES) data from 8 cutaneous squamous cell carcinoma (cSCC) and matched adjacent normal tissue samples^26^. First, we compared the mutations detected by SComatic in epithelial cells using scRNA-seq data with those detected in matched WES data (Methods). For this analysis, we focused on the 9,788,377 positions in the genome across the 8 samples with sufficient coverage in both the scRNA-seq and WES data (Fig. 2d and Methods). In these regions, we detected 266 of the 10,477 (2.4%) mutations found in the WES data, which we considered true positive mutations. Using SComatic, we detected 179 mutations in the scRNA-seq data (Fig. 2d), 78 (44%) of which were also detected in the WES data (Methods). For 49/179 (27%) of the mutations, we found at least 1 read in the WES data supporting the mutated allele, which was however insufficient evidence to call a mutation by our WES analysis pipeline (Methods). Finally, 52/179 (29%) mutations were only detected in the scRNA-seq data. Of these, 38/52 (73%) were detected in sample P7. Interestingly, 59 of the 85 (69%) WES-specific mutations were also detected in P7 only. Mutational signature analysis revealed that 43 (83%) of the mutations only detected in the scRNA-seq data and 70 (82%) of the WES-specific mutations were attributed to single-base substitution (SBS) mutational signatures SBS7a, SBS7b and SBS7d, which are linked with mutagenesis caused by exposure to ultraviolet (UV) radiation, consistent with the expected predominant signature for these samples^26^ (Fig. 2e). In addition, the variant allele fraction (VAF) of the mutations detected in WES and scRNA-seq data were not correlated for P7, unlike for other samples (Supplementary Fig. 3). Therefore, these results suggest that, for sample P7, the lack of sequencing reads in the WES data supporting those mutations detected by SComatic in the scRNA-seq data (and *vice versa*) is likely due to high genetic heterogeneity.

**Figure 2.**
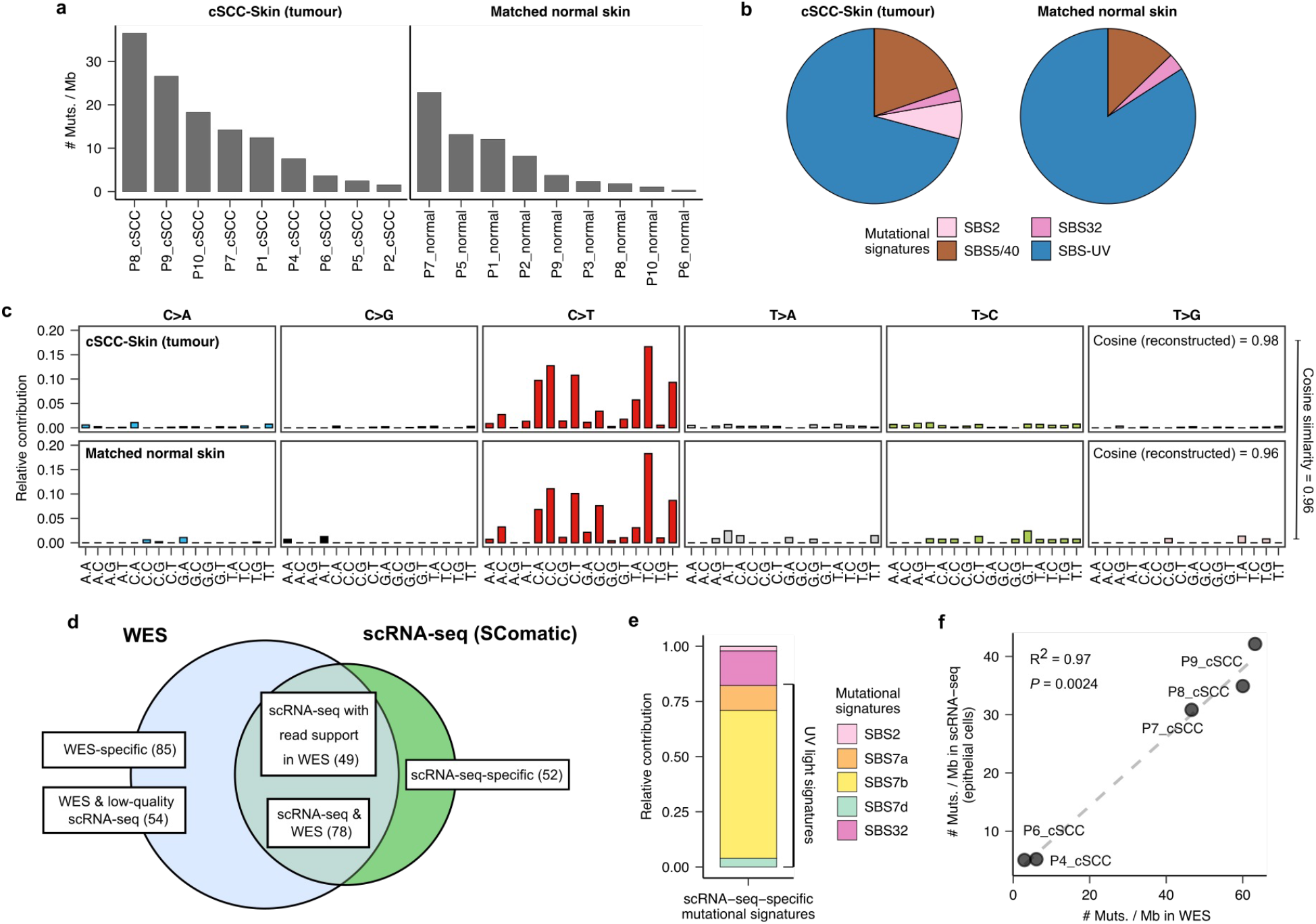
Validation of SComatic using matched scRNA-seq and exome sequencing data. **a**) Mutational burdens for epithelial cells using the somatic SNVs detected by SComatic in cSCC and matched normal skin scRNA-seq data sets. The number of mutations is normalized to account for the variable number of callable sites in each sample. **b**) Fraction of somatic SNVs detected in epithelial cells attributed to COSMIC signatures. SBS signatures associated with ultraviolet radiation (SBS7a,b,c and d) and clock-like mutational processes (SBS5 and SBS40) are collapsed for visualization purposes. **c**) Mutational spectra computed for the mutations detected using SComatic in epithelial cells from cSCC and matched normal skin scRNA-seq data. The cosine similarities between the observed and reconstructed mutational spectra are shown. **d**) Venn diagram showing the overlap of the somatic SNVs detected by SComatic in epithelial cells using scRNA-seq data and exome sequencing data from the cSCC samples. **e**) Decomposition of the mutations detected in scRNA-seq data only (scRNA-seq-specific mutations) into COSMIC signatures. **f**) Correlation between the mutational burdens estimated using the mutations detected in WES and the mutations detected by SComatic in the scRNA-seq data. Only genomic regions with sufficient sequencing depth in both the WES and scRNA-seq data were considered for this analysis.

Next, we applied SComatic to detect somatic mutations across all genomic positions with sufficient coverage in the scRNAseq data (Methods). We detected 810 and 186 SNVs in the tumour and matched normal samples, respectively (Supplementary Table 1), which mapped to 3’-UTR (40%), intronic (27%) and exonic regions (24%) (Supplementary Fig. 4). After normalizing by breadth of coverage (Methods), we estimated an average mutation rate per haploid genome for epithelial cells from the cSCC and normal skin samples of 12.8 and 3.7 mutations per Mb, respectively (note that we report mutational burdens for single cells as mutations per haploid genome because only one allele is usually detected per cell and genomic position). These rates are significantly higher as compared to non-epithelial cells in the data set, which had a median of 0.33 and 0.40 mutations per Mb in tumour and matched normal samples, respectively (*P* < 0.001, Mann-Whitney U-test; Supplementary Fig. 5). Mutational signature analysis attributed 71% and 84% of the mutations detected in epithelial cells from tumour and matched normal skin samples, respectively, to signatures associated with exposure to UV radiation (SBS7a-d; Fig. 2b-c and Methods), consistent with prior DNA sequencing studies of somatic mutations in sun-exposed skin^7,27^. The remaining mutations were mostly attributed to SBS5 and SBS40 signatures (19.6% and 13.4% for the tumour and matched normal samples, respectively), which have been previously identified in non-neoplastic skin samples^7^. The mutation rates computed using the mutations detected using scRNA-seq data for epithelial cells were highly correlated with the rates estimated using the WES data (R^2^ = 0.97, *P* = 0.0024; Fig. 2f and Methods), indicating that SComatic permits the calculation of mutation burdens at cell-type resolution.

Together, these results show a high concordance between the mutations detected in scRNA-seq by SComatic and WES, and highlight that methods for calling mutations in single-cell data based on genotyping mutations previously identified in genome sequencing data are likely to have low sensitivity in samples with high levels of genetic heterogeneity.

### SComatic outperforms existing mutation detection algorithms

Next, we compared the performance of SComatic against top-performing pipelines designed for detecting somatic mutations in scRNA-seq data^22^ using popular variant calling algorithms (VarScan2^28^, SAMtools^29^ and Strelka2^30^). To this aim, we used the matched WES and scRNA-seq data from epithelial cells from 7 out of the 8 cSCC tumours^26^ described above. We excluded patient P7 from this analysis due to the high level of genetic heterogeneity observed between the matched scRNA-seq and WES data (Supplementary Fig. 2). SComatic achieved a sensitivity of 0.59 (95% CI [0.58-0.60]), which was slightly lower than VarScan2 (0.62, 95% CI [0.61-0.63], *P* = 1.86 × 10^−4^), and significantly higher as compared to SAMtools (0.38, 95% CI [0.37-0.39], *P* < 10^−15^). Strelka2 showed a significantly higher sensitivity than SComatic (0.78, 95% CI [0.78-0.79], *P* < 10^−15^; Fig. 3a). However, SComatic outperformed by a large margin all other methods in terms of precision: 0.88 for SComatic (95% CI [0.87-0.89]) *vs* 0.043 for Strelka2 (*P* < 10^−15^, two-sided Student’s *t*-test; Fig. 3a). SComatic also achieved significantly higher F1 score values than other methods (0.71 *vs* < 0.08, respectively; *P* < 10^−15^; Fig. 3a). Notably, we obtained similar differences in performance between methods when also including sample P7 in the benchmarking set (Supplementary Fig. 6).

**Figure 3.**
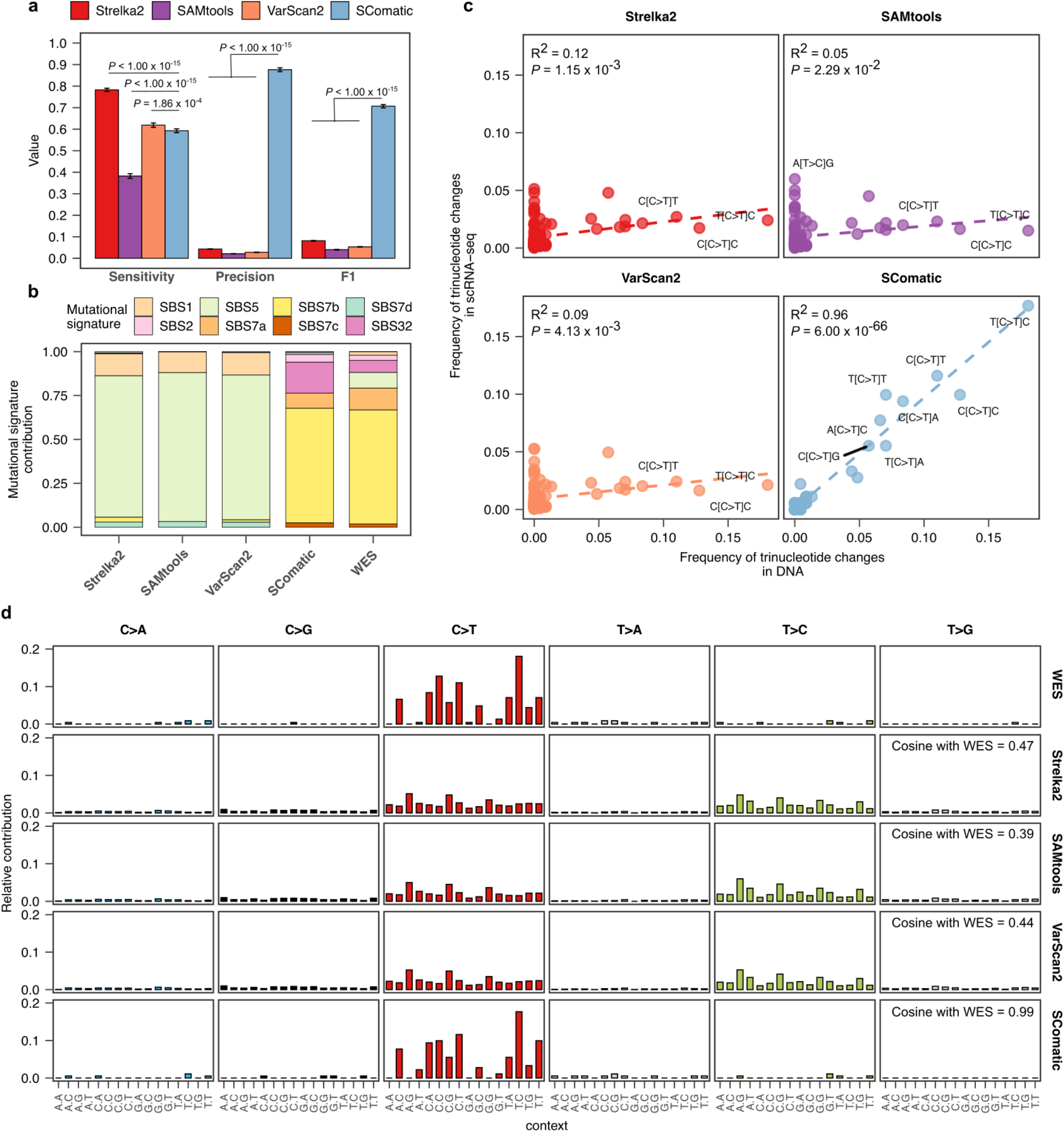
Comparison of the performance of SComatic against other mutation detection methods. **a**) Performance of Strelka2, SAMtools, VarScan2 and SComatic for the detection of somatic mutations in the scRNA-seq data from cSCC samples. The error bars show the 95% bootstrap confidence interval for each statistic computed using 50 bootstrap resamples. **b**) Decomposition into COSMIC signatures of the mutations detected in scRNA-seq data by each algorithm and the mutations detected in WES data. **c**) Correlation between the number of mutations detected in each trinucleotide context using the WES and scRNA-seq data. FDR-adjusted *P* values are shown. **d**) Comparison between the mutational spectra of the mutations detected using WES and scRNA-seq data using each of the algorithms benchmarked. The cosine similarity between the mutational spectra computed using the mutations detected in the scRNAs-seq and the WES data are shown.

To further compare the performance of these algorithms, we performed mutational signature analysis by fitting COSMIC signatures to the observed mutational spectra (Methods). We found that 77% of the mutations detected by SComatic were attributed to signatures SBS7a-d (R^2^ = 0.96 and *P* < 10^−15^, Fig. 3b-c), and the mutational spectrum was highly consistent with the WES data (cosine similarity = 0.99, Fig. 3d). By contrast, the mutations detected by VarScan2, SAMtools and Strelka2 were attributed to signatures SBS1 and SBS5 and were significantly different from the patterns of mutations detected in WES (cosine similarities < 0.47; Fig. 3d). Collectively, these results indicate that existing methods for detecting somatic mutations in scRNA-seq have high false positive rates, whereas SComatic enables the detection of somatic mutations at single-cell resolution at high precision without compromising sensitivity.

### Detection of somatic mutations in samples with high mutational burdens

We next assessed the performance of SComatic to detect somatic mutations in samples characterised by a high mutational burden. To this aim, we applied SComatic to scRNA-seq data from 70 treatment-naïve primary colorectal tumours, including 37 mismatch repair deficient (MMRd) tumours showing microsatellite instability (MSI), and 40 matched normal adjacent colon samples^31,32^. Using SComatic, we called 8,997 somatic SNVs across all samples (7,531 SNVs in MSI, 1,127 in microsatellite stable (MSS), and 339 in the matched normal samples; Supplementary Table 1), most of which mapped to non-coding elements, primarily UTR regions (37%) and introns (27%) (Supplementary Fig. 4). Consistent with previous colorectal cancer genome studies^33,34^, our analysis revealed that epithelial cells in MSI tumours showed a significantly higher mutational burden than epithelial cells from MSS tumours (24.7 vs 8.3 SNVs per Mb, *P* < 1.11 × 10^−12^; two-sided Mann-Whitney U-test) and normal adjacent colon samples (0.51 SNVs per Mb; *P* < 1.77 × 10^−15^). By contrast, the mutational burden for non-epithelial cells was low and comparable between MSI and MSS tumours (0.41 vs 0.52, *P* = 0.06; two-sided Mann-Whitney U-test), as expected for non-malignant cell types (Fig. 4a, Supplementary Fig. 5b). Moreover, the mutational burden estimated by SComatic using scRNA-seq data from epithelial cells in MSI tumours was comparable with that of MMRd tumours estimated using exome-sequencing data from The Cancer Genome Atlas (TCGA)^33,34^ (Fig. 4b; *P* > 0.05; Student’s *t*-test).

**Figure 4.**
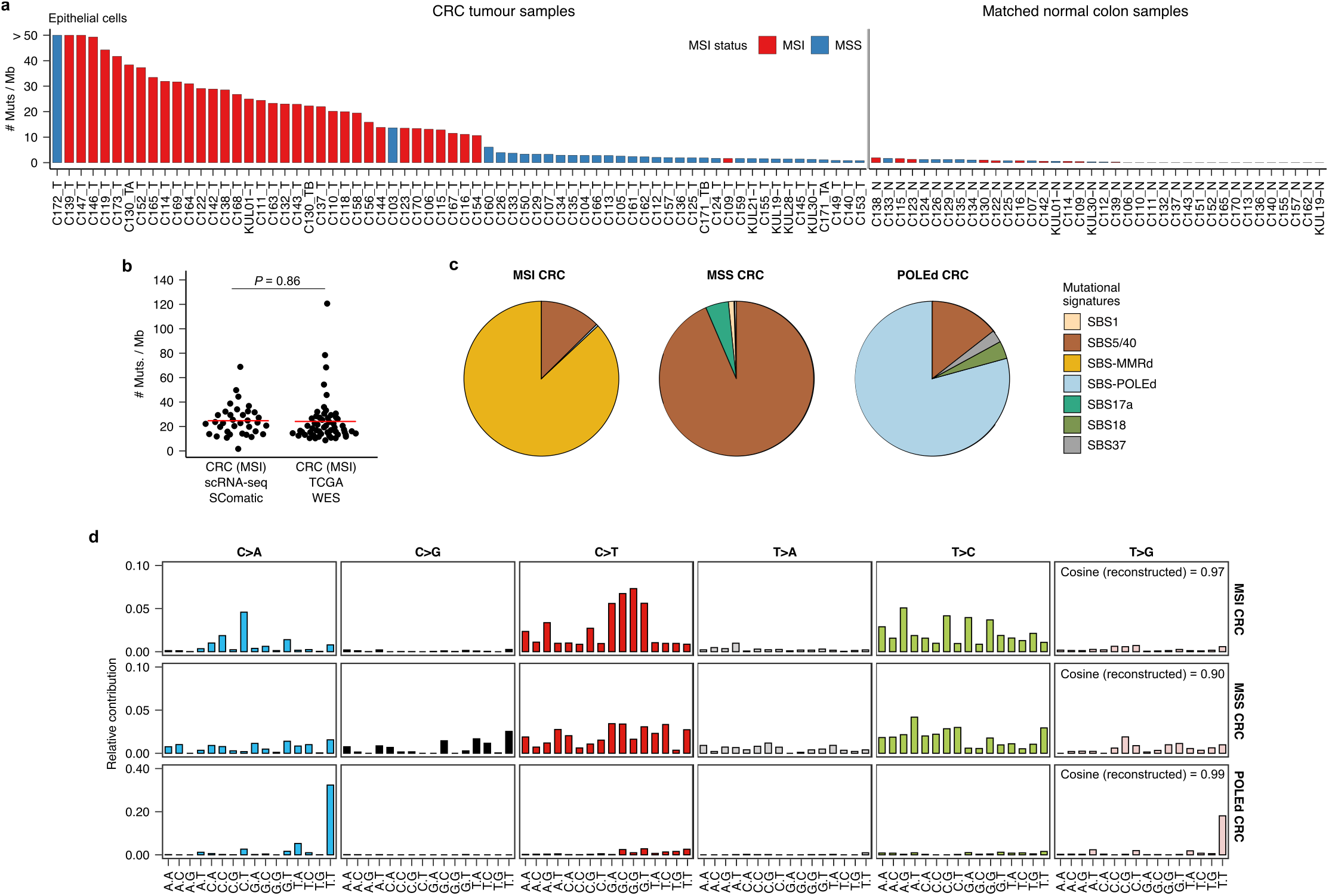
Detection of somatic mutations in scRNA-seq data from colorectal cancer samples. **a)** mutational burden of epithelial cells computed using SComatic. The number of mutations is normalized to the number of callable sites per sample. **b**) Distribution of the mutational burden of epithelial cells from MSI tumours detected using SComatic and the mutational burden of MSI tumours from TCGA computed using WES data. The red horizontal line shows the mean for each group. **c**) Decomposition of the mutational spectra computed using SComatic into COSMIC signatures. Mutational signatures associated with MMRd (SBS6, SBS14, SBS15, SBS21, SBS26 and SBS44), POLE deficiency (SBS10a, SBS10b and SBS28) and clock-like mutational processes (SBS5 and SBS40) are collapsed for visualization purposes. **d**) Trinucleotide context of somatic mutations detected by SComatic using the scRNA-seq data from colorectal cancer samples.

Mutational signature analysis attributed the mutations detected in MSI tumours to SBS signatures associated with MMRd (SBS6, SBS14, SBS15, SBS21, SBS26 and SBS44), SBS5 and SBS40 (Fig. 4c-d; Methods). In one sample (C172), 82.9% of mutations were attributed to signatures SBS10a, SBS10b and SBS28 (Fig. 4a,c,d), suggesting that hypermutation in this sample is driven by *POLE* deficiency^35,36^. In MSS tumours, most mutations were attributed to signatures SBS5 and SBS40, consistent with published compendia of mutational signatures extracted from large cancer genome sequencing studies^36^.

We next compared the mutational burdens estimated by SComatic against VarScan2, SAMtools and Strelka2 using the colorectal cancer scRNA-seq data. As opposed to SComatic, the mutational burdens computed using the mutations detected by the other algorithms were not different between MSI/POLE-deficient and MSS or normal adjacent samples, consistent with the low specificity of existing methodologies for mutation calling using scRNA-seq data (Supplementary Fig. 7).

Together, these results indicate that SComatic permits the identification of the mutational processes operative in hypermutated samples at single-cell resolution without requiring matched genomic sequencing data.

### Detection of mutations using scRNA-seq data from samples with low mutational burdens

We further tested the performance of SComatic to detect mutations in samples with low mutational burdens. To this aim, we applied SComatic to scRNA-seq data for CD34^+^-enriched cells from 5 individuals with myeloproliferative neoplasms (MPN), a type of blood cancer caused by the clonal expansion of a single hematopoietic stem cell (HSC)^8^. We detected an average of 0.12 mutations per Mb per haploid genome, which primarily mapped to intronic regions (62%, Supplementary Figure 4). Mutational signature analysis revealed that 96% of the mutations detected by SComatic were attributed to signatures SBS5 and SBS40 (Fig. 5a-b), consistent with single-cell whole-genome sequencing (WGS) studies of HSCs from healthy donors^6,37^ and MPN patients^8,38^. In addition, we found a positive correlation between the average mutation rate of HSCs estimated by SComatic and the patients’ age at the time of sampling (Pearson’s r = 0.79; *P* = 0.09, Fig. 5c), in agreement with previous studies^8^. Altogether, these results show that SComatic accurately detects mutational burdens and signatures in samples with low mutational burdens.

**Figure 5.**
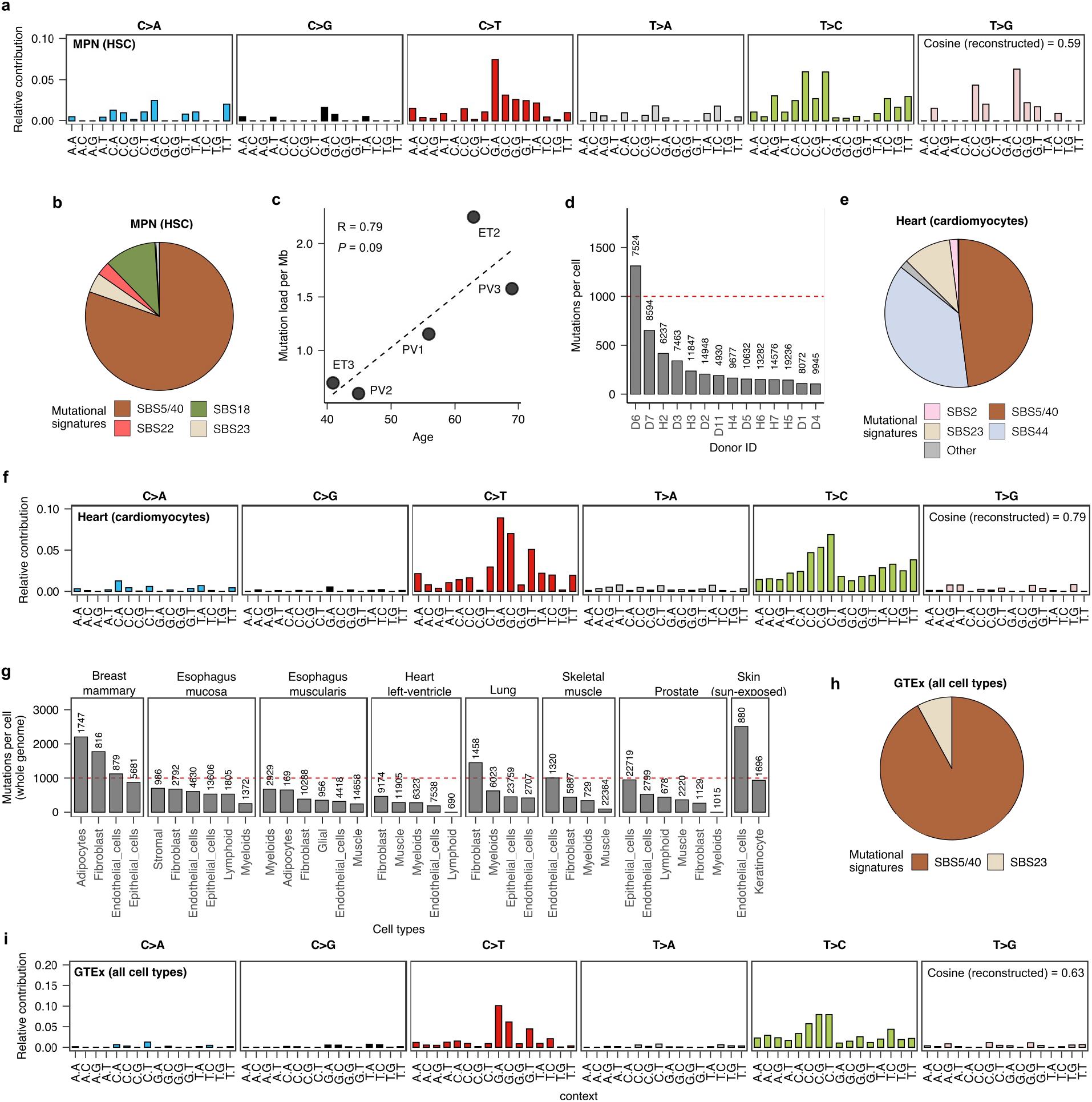
Detection of somatic mutations in samples with a low tumour mutational burden. **a)** Trinucleotide context of somatic mutations detected in hematopoietic stem cells (HSC) from MPN patients. **b**) Decomposition of the somatic mutations detected in HSCs from MPN patients into COSMIC signatures. **c**) Correlation between the mutational burden of HSCs estimated using SComatic and the age of patients at the time of sampling (Pearson’s r = 0.79; *P* = 0.09). **d**) Average number of mutations detected per cell and genome in cardiomyocytes from the heart cell atlas across donors. **e**) Decomposition of the mutations detected in cardiomyocytes into COSMIC signatures. **f**) Trinucleotide context of mutations detected in cardiomyocytes from the heart cell atlas. **g**) Average mutational burden of individual cells across the tissues included in the GTEx scRNA-seq dataset. The number on top of the bars indicates the number of cells per cell type. **h**) Decomposition of the mutations detected across all cells from the GTEx data set into COSMIC signatures. **i**) Trinucleotide context of mutations detected across all single cells from the GTEx data set. The numbers on top of the bars in **d** and **g** indicate the number of cells per cell type analysed.

To further test whether SComatic can be used for the analysis of somatic mutations in samples with high levels of genetic heterogeneity (e.g., polyclonal tissues) and in differentiated cells, we next analysed 10X scRNA-seq data from 78 samples obtained from 6 heart regions across 14 donors^39^. We detected a total of 2,132 somatic SNVs (Supplementary Table 1), 78% of which mapped to intronic regions (Supplementary Fig. 4). By extrapolating to the entire genome, we estimated an average mutation rate per haploid genome of 302 mutations for cardiomyocytes (range 92-1,284; Fig. 5d), which was significantly lower than the mutation rates estimated for adipocytes (1,179 SNVs per cell and haploid genome) and smooth muscle cells (581; Supplementary Fig. 8a). Mutational signature analysis revealed that 46.7% of these mutations were attributed to SBS5 and SBS40 (Fig. 5e,f). In addition, 35.4% of mutations were attributed to SBS44, consistent with a recent study of somatic mutagenesis in human cardiomyocytes using single-cell genome sequencing^40^. The mutational burdens for cardiomyocytes estimated by SComatic were comparable to those estimated using single-cell WGS data^40^ (Supplementary Fig. 9; *P* = 0.08; two-sided Wilcoxon’s rank test).

Next, we applied SComatic to 24 scRNA-seq data sets from 8 non-neoplastic tissues across 15 human donors generated by the GTEx consortium^41^. We found a total of 524 SNVs and estimated an average mutation load of 598 mutations per cell and haploid genome (Fig. 5g, Supplementary Fig. 8b and Methods). As observed in the heart cell atlas, adipocytes had the highest mutation burdens (1,430 mutations per cell and haploid genome), whereas muscle cells showed the lowest burdens (251; Supplementary Fig. 8b). As observed in other polyclonal tissues^7^, mutational signature analysis revealed that most of these mutations were attributed to the mutational signatures SBS5 and SBS40 (92.1 %, Fig. 5h,i). Together, these results suggest that SComatic permits the study of the patterns and rates of mutations in polyclonal tissues.

### Performance of SComatic on single-cell ATAC-seq data sets

Next, we applied SComatic to detect somatic mutations using sciATAC-seq data generated for 459,056 cells from 66 samples spanning 24 non-neoplastic tissues^42^. SComatic detected a total of 389 somatic SNVs (Supplementary Table 1). The distribution of mutations was different as compared to scRNA-seq data sets, as most mutations mapped to intergenic (32%), promoter (19%), and intronic regions (18%) (Supplementary Fig. 4). We found low single-cell mutational burdens with an average load of 300 mutations per cell and haploid genome, with ductal cells showing the highest rates (933 per haploid genome), and skeletal myocytes (9 mutations) and follicular cells (0 mutations) the lowest burdens (Supplementary Figs. 10a-c). As observed in other polyclonal tissues, 99% of the SNVs were attributed to SBS5 and SBS40 (Supplementary Fig. 10b,c). Importantly, the genome-wide mutation rates were comparable for cell types represented in scRNA-seq and sciATAC-seq data sets, indicating that SComatic permits the estimation of mutation rates across different single-cell profiling assays (Supplementary Fig. 11).

### Patterns of clonality at cell-type resolution

Motivated by the importance of clonal mosaicism to somatic evolution and disease^2,43^, we next assessed whether the single-cell resolution provided by SComatic permits analysis of the patterns of clonality across cell types. To this aim, we computed the fraction of mutant cells per cell type across the single-cell data sets analysed (Supplementary Table 1, Supplementary Fig. 12 and Methods). We detected clonal mutations in epithelial cells from the cSCC samples, but not in epithelial cells from non-neoplastic skin samples, consistent with the high level of polyclonally in normal skin (Supplementary Fig. 12a,b). The clonality of mutations in epithelial cells in both MSI and MSS colorectal samples spanned a dynamic range of values, as expected for tumours harbouring both clonal and subclonal mutations (Supplementary Fig. 12c,d). The mutations detected in non-neoplastic cell types from both cancer and non-neoplastic samples showed overall low (<0.2) mutant cell fractions, in agreement with genome sequencing studies of non-neoplastic tissue samples^7^ (Supplementary Fig. 12d-f). Together, these results show that SComatic permits the study of clonality patterns of both cancer and non-neoplastic cell types.

### *De novo* mutational signature analysis

Clustering of samples based on the cosine similarity of mutational spectra revealed groups consistent with the relative activity of known mutational processes quantified though refitting of COSMIC mutational signatures (Supplementary Fig. 13). Thus, we sought to determine whether the mutations detected by SComatic permit the identification of mutational processes using *de novo* mutational signature extraction. Decomposition of the mutations identified in epithelial cells from hypermutated colorectal cancer samples using COSMIC signatures revealed a strong contribution of signatures associated with POLE and MMRd. By contrast, the signatures extracted from epithelial cells in MSS tumours showed strong contributions of SBS5 and SBS40, consistent with the mutational processes expected for these tumours (cosine similarities > 0.96, Supplementary Fig. 14). We identified two signatures in cSCC samples, one of which showed a cosine similarity >0.98 when decomposed into the COSMIC signatures attributed to UV-light mutagenesis (SBS7a, SBS7b and SBS7c), and the other was decomposed into a combination of signatures (SBS5 and SBS40), in agreement with the WES data (cosine similarity = 0.7, Supplementary Fig. 14). Despite the limited number of mutations and samples available for analysis, the signatures extracted from the mutations detected in non-neoplastic samples from GTEx and the heart cell atlas were decomposed into SBS5 and SBS40 (cosine similarity > 0.36; Supplementary Fig. 14), which is consistent with the mutational signatures identified in WGS studies of non-neoplastic samples^7^. The signatures detected in cardiomyocytes showed a strong contribution of SBS44, which is related to MMRd and recently reported in a recent study of cardiomyocytes using single-cell WGS^40^. Together, these results indicate that SComatic permits *de novo* mutational signature analysis using mutations detected in single-cell data.

## Discussion

Here, we show that SComatic permits *de novo* detection of somatic SNVs at single-cell resolution. In contrast to existing methods relying on genotyping sites known to be mutated in the sample under study, SComatic detects somatic SNVs in single-cell data sets directly without requiring matched bulk or single-cell DNA sequencing data. This is particularly relevant to study somatic mutagenesis in cell types and samples that cannot be reliably analysed using existing single-cell genomics methods, such as differentiated cells and polyclonal tissues showing high levels of genetic heterogeneity^5,7^. Critically, we show that SComatic vastly outperforms existing pipelines for the detection of somatic SNVs in single cell data sets, which allows the identification of mutational processes in both cancer and non-neoplastic cells, including those from differentiated cells and polyclonal tissues in which mutations cannot be reliably studied using current experimental or computational approaches.

Despite its higher performance as compared to existing tools, we note that SComatic is limited by the sparsity and low sequencing depth of current single-cell sequencing assays. As single-cell methods improve, SComatic will allow to derive further insights from single-cell sequencing data sets, such as phylogenetic analysis, identification of driver mutations in cancer and non-neoplastic cells, and the study of clonal mosaicism, including the estimation of mutations under positive selection driving clonal expansions. Although we have previously shown that somatic mutations can be detected in off-target regions, such as introns^44^, only a small fraction of the genome has sufficient sequencing coverage to be amenable to mutation detection. Therefore, other methodologies are required to study the rates, patterns, and selection of mutations in those regions missed by scRNA-seq and ATAC-seq or overlapping known RNA editing sites. In addition, SComatic relies on predefined cell type annotations using e.g., marker genes or gene expression clustering. Therefore, the quality of the mutations identified is contingent on reliable cell type annotations, which can be challenging in cases in which clonally unrelated cells cannot be easily distinguished using gene expression data alone^8,44^. Finally, we applied SComatic to study the patterns of clonality and mutation rates in clonal and polyclonal tissues. Although the cell-type mutation rates we estimate are comparable across assays, we note that the bias introduced by allele-specific expression, polyploidization, and limited sequencing depth might affect the burden or clonality estimates for other data sets.

Overall, SComatic opens the possibility to study somatic mutagenesis using single-cell data sets generated for human samples under the auspices of large-scale initiatives, such as the Human Cell Atlas or the Human Tumour Atlas Network^45,46^, as well as the analysis of mutational burdens and processes in other organisms.

## Methods

### Processing of single-cell data sets

Single-cell RNA-seq data from cancer and non-neoplastic samples were downloaded in fastq format and processed uniformly. Specifically, raw sequencing reads were aligned to the GRCh38 build of the human reference genome using Cell Ranger^47^ version 6.0.1 and default parameter values to generate alignment files in Binary Alignment Map (BAM) format and count matrices. Cell type annotations were downloaded from the original publications from which the data were downloaded (Supplementary Table 1). Cell annotations were used to assign sequencing reads to individual cells. Single cells without cell type annotations were discarded. Raw sciATAC-seq reads were mapped to the GRCh38 build of the human reference genome using BWA-MEM v0.7.17-r1188^48^. Aligned sequencing reads in BAM format were then processed following the Genome Analysis Toolkit (GATK) v4.1.8.0 Best Practices workflow to remove duplicates and recalibrate base quality scores^49^.

### Detection of somatic mutations in single-cell data sets using SComatic

SComatic consists of the following steps:

#### Processing of alignment files

First, the BAM file containing the sequencing reads for all cell types in a sample is split into cell-type-specific BAM files using precomputed cell type annotations. To this aim, sequencing reads are assigned to individual cells using molecular barcodes (tag “CB” in BAM files processed using Cell Ranger). Before identifying candidate mutation sites, reads with a mapping quality lower than 255 (or 30 for sciATAC-seq data) or with more than 5 mismatches are filtered out. In addition, to ignore sequencing artefacts enriched in terminal ends of the reads or adapter sequences not properly trimmed, the base quality for the first 5 bases at the 3’ and 5’ ends of each read is set to 0^50^.

#### Collecting base count information

Next, the count of each base in each cell type for every position in the genome is recorded in a base count matrix indexed by cell types and genomic coordinates using the pileup functionality from the Pysam module^51^. For this analysis, a minimum base quality of 30 is required, and only sites with a sequencing depth of 5 reads across at least 2 cell types are considered. Sites overlapping RNA editing sites are removed^52,53^. In addition, sites mapping to polymorphisms in the gnomAD^25^ database version v2.0.1 with a population frequency greater than 1% are removed.

#### Detecting potential somatic SNVs

To distinguish technical artefacts, such as recurrent sequencing or mapping errors, from true somatic mutations, SComatic models the background error rate using a Beta-binomial distribution. Specifically, non-reference allele counts at homozygous reference sites are modelled using a binomial distribution with parameter P (error rate), which is a random variable that follows a Beta distribution with parameters α and β^50^. To infer the parameter values, SComatic uses base count information for 1 million sites in the genome randomly selected from a panel of unrelated non-neoplastic samples generated using the same sequencing technology. Next, for each site in the genome and cell type, the Beta-binomial distribution is used to test whether the non-reference allele counts are significantly higher than expected given the background error rate, and thus, considered as a potential somatic mutation. Candidate somatic mutations are required to be present in only cells from the candidate cell type. To test this, SComatic requires that the Beta-binomial test is not significant when applied to all other cell types independently and when applied to the base counts aggregated across all other cell types. The threshold for statistical significance for the Beta-binomial is set to 0.001.

#### Filtering out recurrent artefacts

Due to the enrichment of artefacts in repetitive regions (Supplementary Fig. 1) and the high error rate of Illumina sequencers at homopolymer tracts^54^, mutations mapping to or within 4bp of mononucleotide tracts are removed. Finally, mutations mapping less than 5bp apart from each other are filtered out, except for doublet base substitutions (DBS) dinucleotide changes previously reported to be generated by specific mutational processes, such as CC>TT mutations associated with UV-light-induced mutagenesis in skin (COSMIC signature DBS1) or characteristic DBS peaks observed in colorectal cancers (COSMIC signatures DBS2,3,4,6,7,8,10 and 11).^36^

In addition, SComatic generates a ‘Panel of Normals’ to discount positions affected by recurrent artefacts (sites with non-reference allele counts significantly higher than the background error rate modelled with the Beta-binomial distribution). To this aim, SComatic uses a large collection of non-neoplastic datasets to assess the frequency of non-reference allele counts at each genomic site in the genome. This analysis serves to filter out candidate mutations mapping to regions of the genome prone to sequencing or mapping artefacts, germline variants missed by other filters, and candidate mutations found in at least 2 unrelated samples, which are considered to be germline polymorphisms.

#### Calling somatic mutations

Finally, to make a mutation call, SComatic requires mutations to be supported at least 3 reads from at least 2 cells from the same cell type. To tune this parameter, we performed mutational signature analysis on subsets of mutations defined based on the number of cells harbouring each mutation. For this analysis, we focused on the somatic mutations detected by SComatic in epithelial cells from MSI tumours. Our analysis revealed that the mutational spectra and mutational signature contributions were consistent across subsets of mutations present in 2 or more cells (Supplementary Fig. 2), indicating that requiring mutations to be present in at least 2 cells to make a call is adequate to detect true somatic mutations

### Estimation of mutational burdens

To compute the mutational burden at the cell type level, we divided the total number of somatic mutations detected in each cell type by the total number of callable sites across all cells of the same type (Supplementary Fig. 15). Cell types with less than 500,000 callable sites were not included in this analysis. To estimate single-cell mutational burdens, we divided the number of mutations detected in each unique cell by the number of sites with a sequencing depth of at least 1 read and within the set of callable sites across all cells of the same type. We only considered the autosomes for computing mutational burdens. The sensitivity of single-cell assays to detect both alleles is low due to limited sequencing depth and allele-specific expression^17^. That is, we only detect one read per cell for most genomic position in the genome. Thus, our estimated mutational burdens for single cells mostly reflect the mutational burdens per haploid genome. We decided to report mutational burdens per haploid genome instead of correcting for ploidy because ploidy information for single cells was not available for the data sets analysed. We could not assume that all cells are diploid as the data sets analysed contained cell types, such as cancer cells and cardiomyocytes, that often undergo polyploidization.

### Mutational signature analysis

Mutational signature analysis was performed using the R package MutationalPatterns^55^ and the COSMIC Mutational Signatures catalogue version 3^36^. We used the function *fit_to_signatures* with default parameter values to estimate the contribution of each mutational process to the mutational spectrum observed in each sample. To account for differences in the frequency of each of the 96 trinucleotide contexts in which mutations can be detected between the whole genome and the regions profiled using scRNA-seq or scATAC-seq, we normalised the frequency of mutations detected at each trinucleotide context. To this aim, we first computed the frequency of each trinucleotide context in the human genome using the function *get_trinuc_norm* from the R package SigMA (https://github.com/parklab/SigMA). Next, for each single-cell data set we estimated the frequency of each trinucleotide context across callable regions using a custom Python script, *TrinucleotideContextBackground*.*py*, which is provided as part of SComatic. To normalize the mutational spectra detected in each single-cell data set to the frequency of each trinucleotide in the whole genome, we divided the fraction of mutations detected at each trinucleotide context by the frequency of such context in the whole genome relative to its frequency in the single-cell data set being analysed.

For fitting COSMIC signatures, we only used the mutational processes known to be operative in each sample type analysed^7,36^: (1) SBS1, SBS5, SBS6, SBS10a, SBS10b, SBS14, SBS15, SBS17a, SBS17b, SBS18, SBS21, SBS26, SBS28, SBS37, SBS40 and SBS44 for colorectal cancer samples; (2) SBS1, SBS2, SBS5, SBS7a, SBS7b, SBS7c, SBS7d, SBS13, SBS32 and SBS40 for skin squamous cell carcinoma samples; (3) SBS1, SBS2, SBS4, SBS5, SBS7a, SBS7b, SBS13, SBS16, SBS17b, SBS18, SBS22, SBS23, SBS32, SBS40, SBS41 and SBS88 for MPNs and non-neoplastic samples. We also included SBS6, SBS8, SBS19, SBS32, SBS35, SBS39, and SBS44 when analysing heart samples^40^. The goodness of fit was determined by computing the cosine similarity between the observed and the reconstructed mutational spectra using the estimated signature contributions.

*De novo* mutational signature extraction was performed using non-negative matrix factorization (NMF) as implemented in the R package *MutationalPatterns* using somatic SNVs detected in each of the following sample groups: epithelial cells from MSI and POLE-deficient colorectal cancer samples, epithelial cells from MSS colorectal cancer samples, epithelial cells from cSCC and matched normal skin samples, cardiomyocytes from the heart cell atlas, and all cell types from the GTEx dataset. The extracted signatures were decomposed into COSMIC v3 signatures using the *fit_to_signatures* function after normalizing them to the trinucleotide frequencies of the whole genome. The goodness of fit of the decomposition of d*e novo* signatures was estimated by computing the cosine similarity between the extracted mutational signature and the mutational spectrum reconstructed based on the estimated COSMIC signature contributions.

### Whole-exome sequencing data analysis

Raw sequencing reads were mapped to the GRCh38 build of the human reference genome using BWA-MEM^29^ (version 0.7.17-r1188). Aligned sequencing reads in BAM format were processed to remove duplicates and recalibrate base quality scores following the GATK (version 4.1.8.0) Best Practices workflow^56,57^. Point mutations were detected using Strelka2^30^ (version 2.9.10) and MuSE^58^ (version 1.0rc) using default parameter values and the matched normal samples as germline controls. For benchmarking purposes, we only considered those somatic mutations detected by both algorithms.

### Comparison of mutations detected in scRNA-seq and WES data

To compare the mutations detected using matched WES and scRNA-seq data, we computed the base counts for all positions in the genome using the WES data. For this analysis, we only focused on regions with a coverage of at least 50x in the WES data from the cancer sample and 10x in the matched normal sample. In the case of the scRNA-seq data, we only interrogated regions with a sequencing depth of at least 10 reads in the epithelial cells, and with a depth of 5 reads in at least 2 additional cell types. Only regions that passed these filtering criteria for the scRNA-seq and WES data were considered for benchmarking purposes.

As we considered the WES data as the baseline for comparison, we categorized the mutations as: (1) true negatives: non-mutated sites; (2) WES-specific mutations: mutations detected in the WES but not in scRNA-seq data; (3) scRNA-seq-specific: mutations detected in the scRNA-seq data with no reads supporting the mutant allele in the WES data; (4) low-confidence true positives: mutations detected in the scRNA-seq data with at least one read supporting the alternative allele and no reads supporting any other alternative allele in WES, but not called by our WES mutation detection pipeline; (5) true positives: mutations detected in both the scRNA-seq and WES data; and (6) WES and low-quality scRNA-seq: somatic mutations detected in WES but filtered out by SComatic. To compute performance metrics, we estimated the sensitivity, precision and F1-score values for each algorithm using 50 bootstrap resamples. We then compared the performances between callers using the Student’s *t*-test correcting for multiple hypothesis testing using the FDR method.

## Data availability

The raw WES and scRNA-seq data for the skin squamous cell carcinoma and matched normal samples are available at the Gene Expression Omnibus (GEO) database under the accession number GSE144240. The raw scRNA-seq data from myeloproliferative neoplasms and colorectal cancer patients are available through controlled access application via dbGaP under dbGaP Study Accession numbers phs002308.v1.p1 and phs002407.v1.p1, respectively. The cell type annotations for the colorectal cancer data set^31^ are available at GEO database under the accession number GSE178341. Raw sequencing data and cell type annotations for 6 additional colorectal cancer patients^32^ included in this study are available at GEO database under the accession number GSE144735. The cell type annotations for the MPN data set were obtained from our previous study^8^. The raw scRNA-seq data and cell type annotations for the human heart cell atlas^39^ were downloaded from the Human Cell Atlas Data Portal (https://data.humancellatlas.org/). The raw single-cell ATAC-seq data and cell type annotations are available at GEO database under the accession number GSE184462. The raw sequence data from GTEx samples are available at the Analysis Visualization and Informatics Lab-space (AnVIL; https://anvil.terra.bio/#workspaces/anvil-datastorage/AnVIL_GTEx_V9_hg38) and can be downloaded through controlled data access application via dbGaP under Study Accession number: phs000424.

## Code availability

SComatic is available at: https://github.com/cortes-ciriano-lab/SComatic.

## Author contributions

I.C.-C. designed and supervised the study. F.M. performed analyses, generated the figures, and implemented SComatic with input from I.C.-C. I.C.-C. and F.M. wrote the manuscript with input from all authors. R.L., R.R., T.M. and S.H. contributed to discussing the results. All authors read and approved the final version of the manuscript.

## Supplementary Figures

**Supplementary Figure 1.**
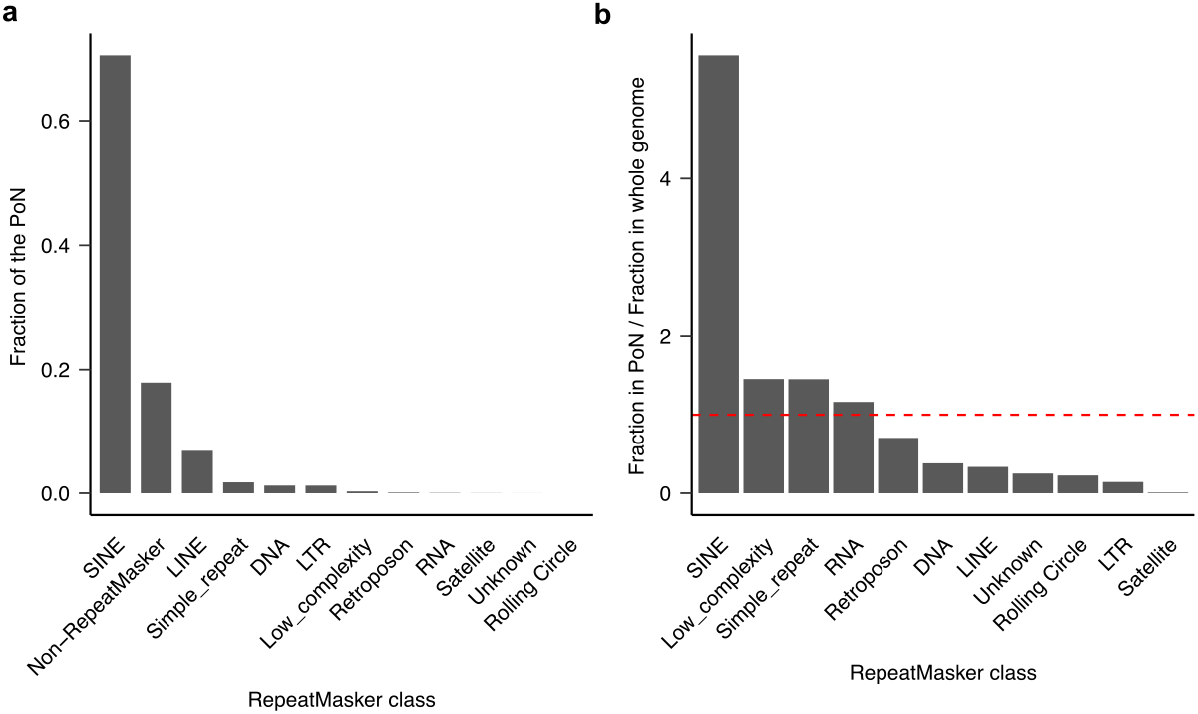
Recurrent artefacts are enriched in repetitive elements. **a**) Genomic distribution of artefactual sites included in the “Panel of Normals” (PoN) generated using scRNA-seq data across different types of repetitive elements. **b**) Enrichment of artefactual sites in repetitive element classes. The dashed red line indicates no enrichment or depletion.

**Supplementary Figure 2.**
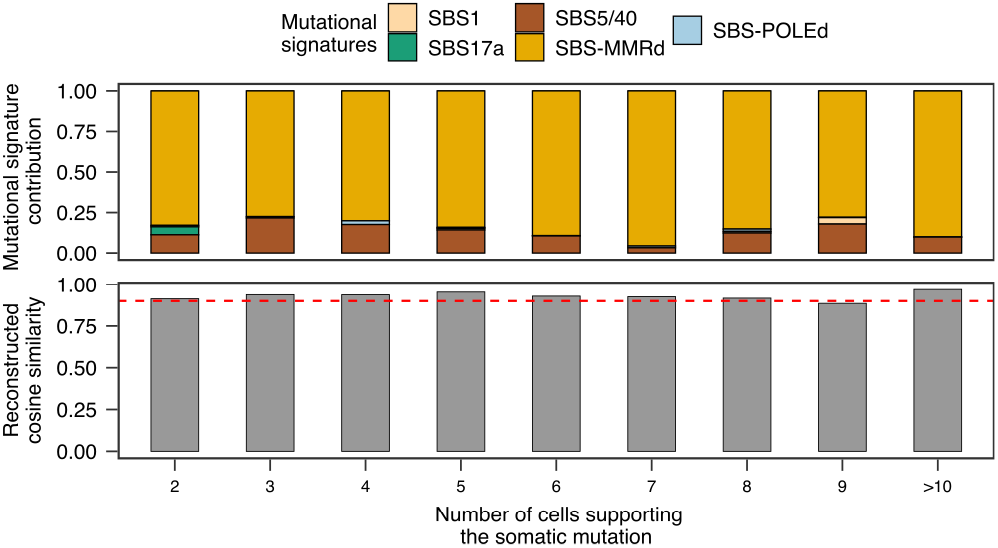
Mutational signature analysis of the somatic mutations detected across an increasingly larger number of cells. Decomposition into COSMIC signatures of the somatic mutations detected in the MSI colorectal cancer data sets across increasingly higher cut-off values for the number of cells required to harbour a mutation to make a call. Overall, the contribution of mutational signatures associated with MMRd is constant across increasingly stringent cut-off values, indicating that requiring mutations to be detected in at least 2 cells to make a call is adequate to discover true somatic mutations. Mutational signatures associated with MMRd (SBS6, SBS14, SBS15, SBS21, SBS26 and SBS44), POLE-deficiency (SBS10a, SBS10b and SBS28) and clock-like mutational processes (SBS5 and SBS40) are collapsed for visualization purposes.

**Supplementary Figure 3.**
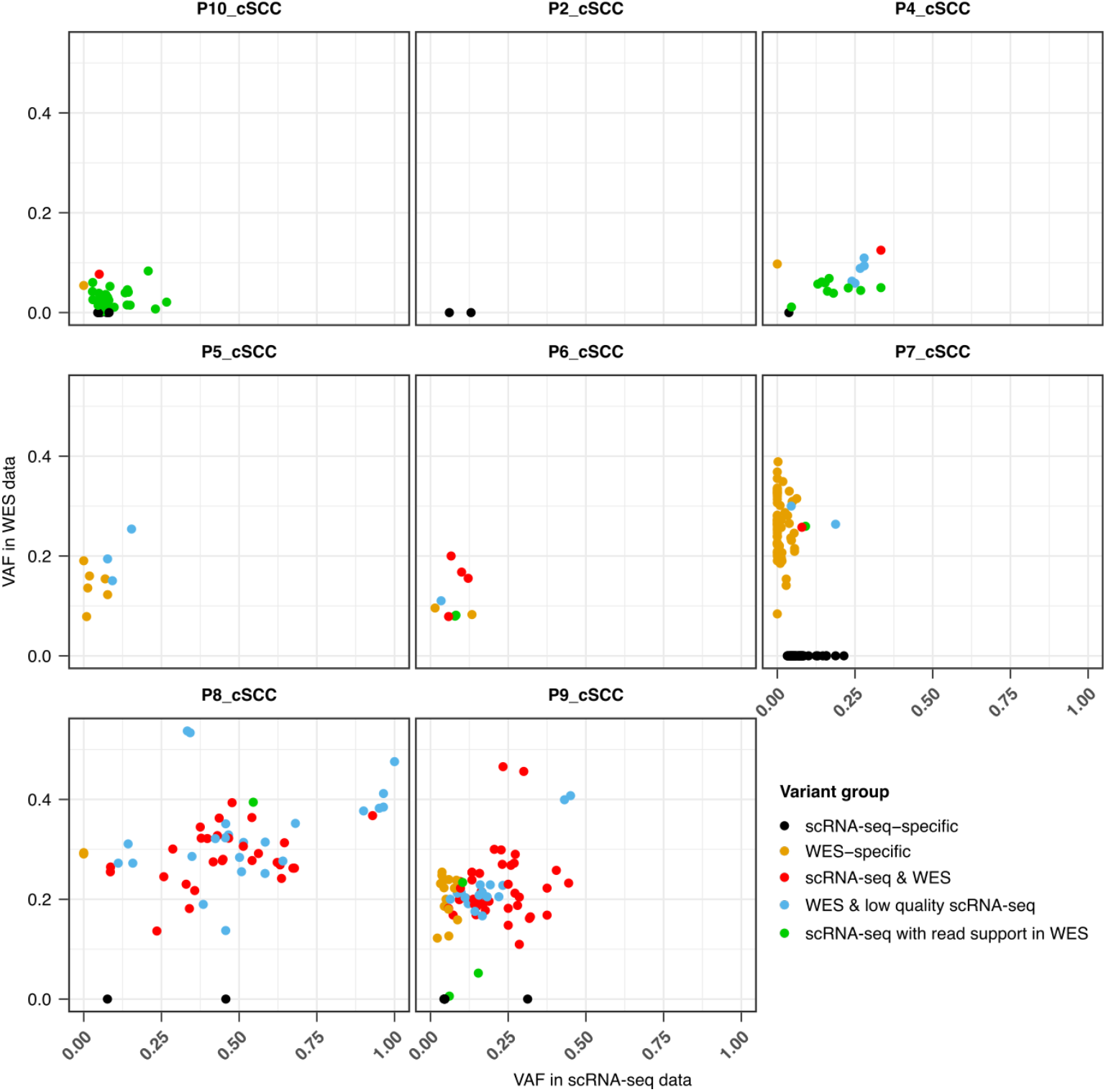
Comparison of the variant allele fraction (VAF) of mutations detected in WES data and scRNA-seq data from epithelial cells using SComatic.

**Supplementary Figure 4.**
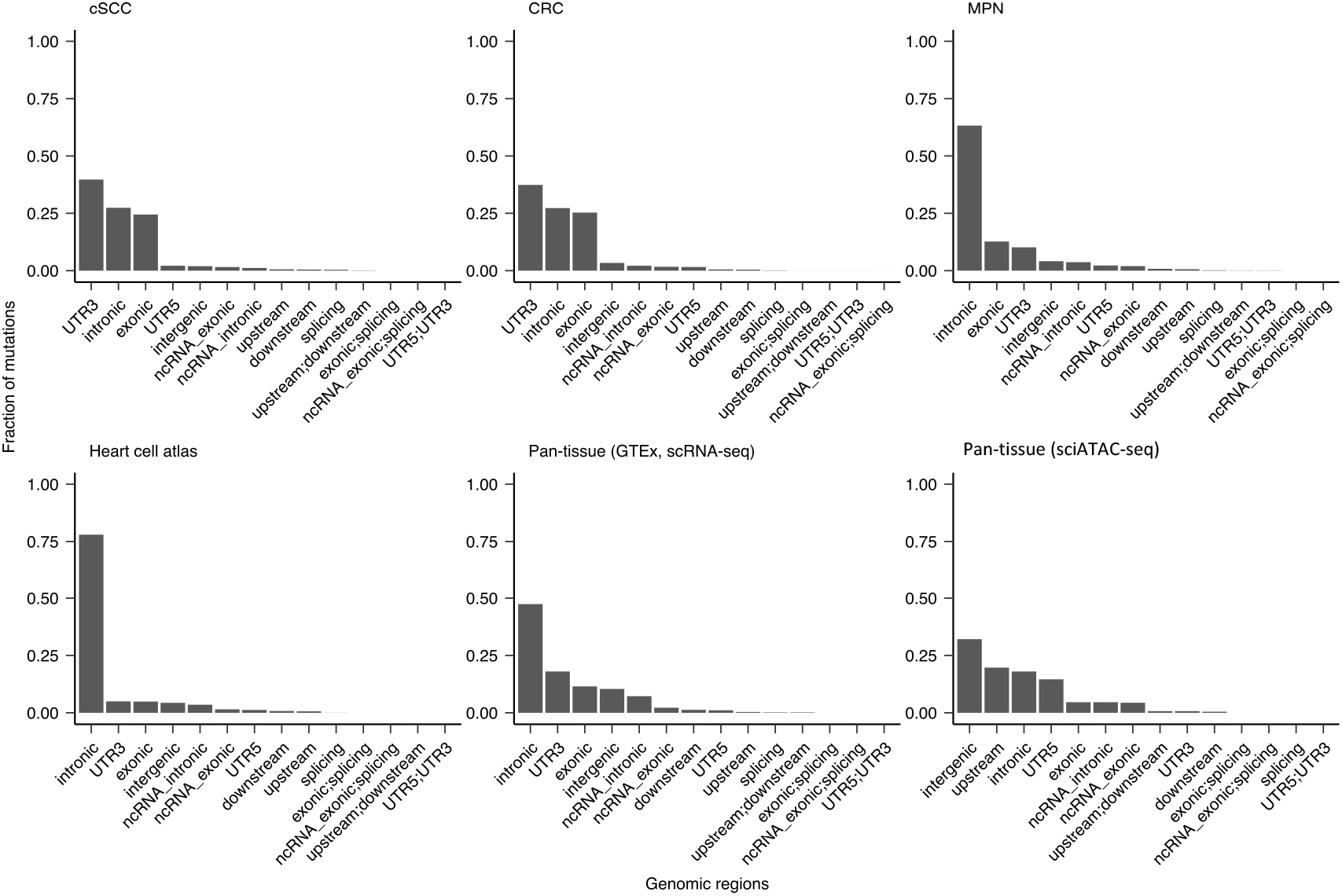
Genomic distribution of somatic mutations detected by SComatic in single-cell data sets. Distribution of somatic mutations across genomic regions.

**Supplementary Figure 5.**
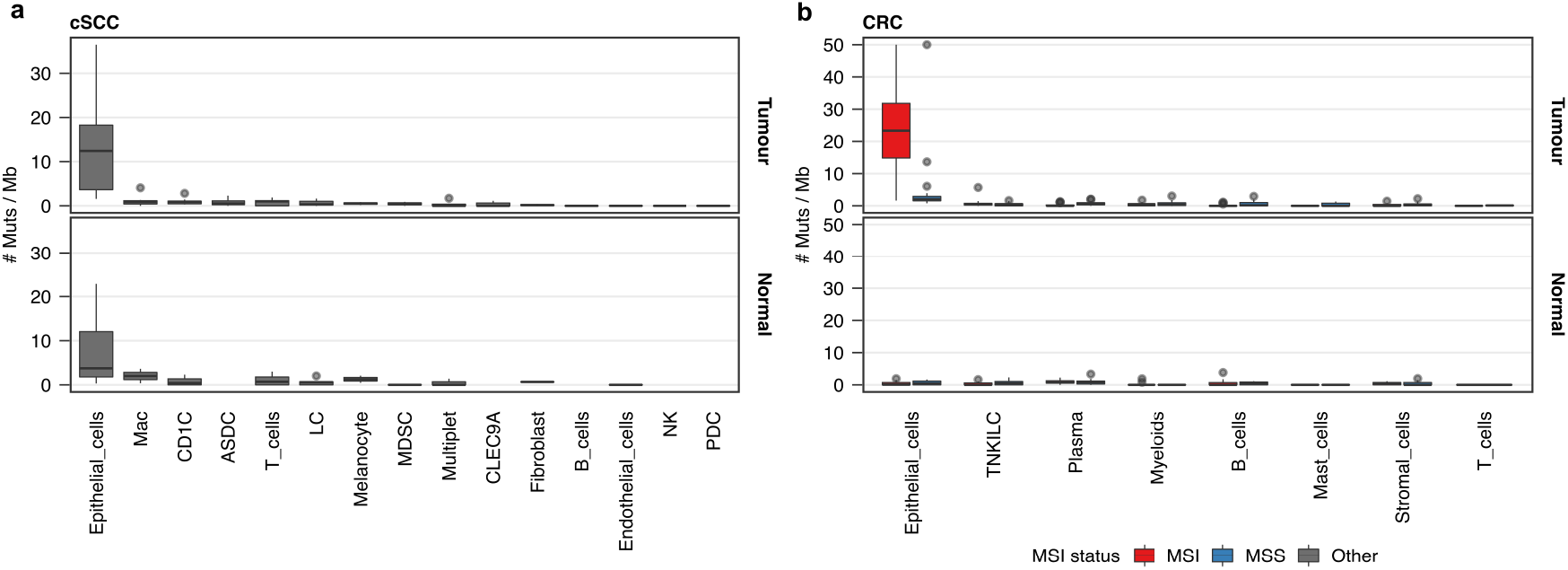
Mutational burden across cell types. **a**) Mutational burden across cell types detected in the scRNA-seq data from cSCC and matched normal skin samples. **b**) Mutational burden across cell types detected in the colorectal cancer (CRC) and matched normal colon samples. Mac: macrophages; ASDC: AXL+SIGLEC6+ dendritic cells; LC: Langerhans cells; MDSC: myeloid-derived suppressor cells; PDC: Plasmacytoid dendritic cells; TNKILC: T-cells, natural killer cells, Innate lymphoid cells. Box plots show median, first and third quartiles (boxes), and the whiskers encompass observations within a distance of 1.5× the interquartile range from the first and third quartiles.

**Supplementary Figure 6.**
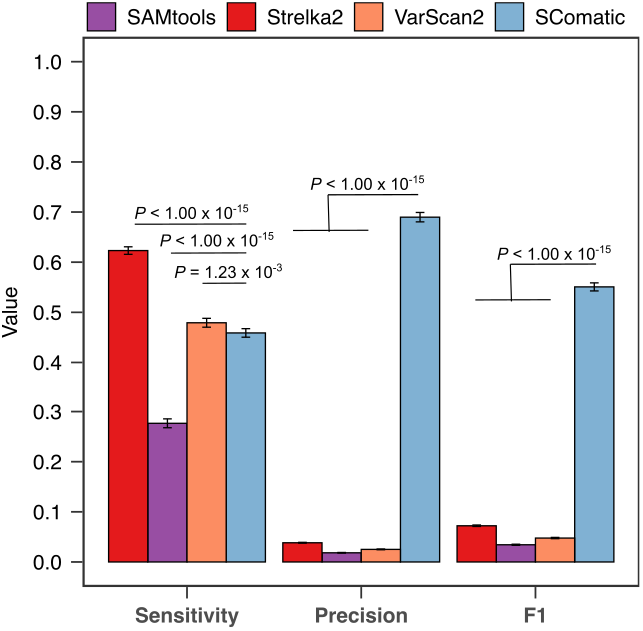
Comparison of the performance of SComatic against other mutation detection methods including sample P7. Performance of Strelka2, SAMtools, VarScan2 and SComatic for the detection of somatic mutations in scRNA-seq data from cSCC samples (including sample P7). The error bars show the 95% bootstrap confidence interval for each statistic computed using 50 bootstrap resamples.

**Supplementary Figure 7.**
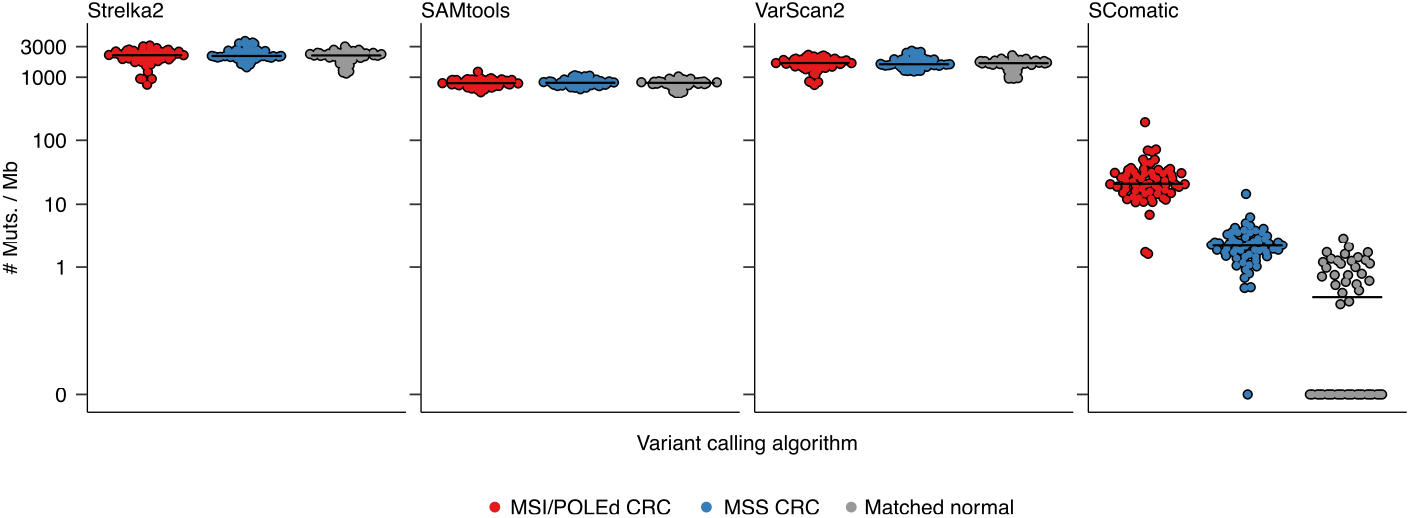
Comparison of the mutational burden of epithelial cells computed using the mutations detected by Strelka2, SAMtools, VarScan2 and SComatic using the scRNA-seq data from colorectal cancers. Each dot represents a sample, and the black horizontal line shows the median for each group.

**Supplementary Figure 8.**
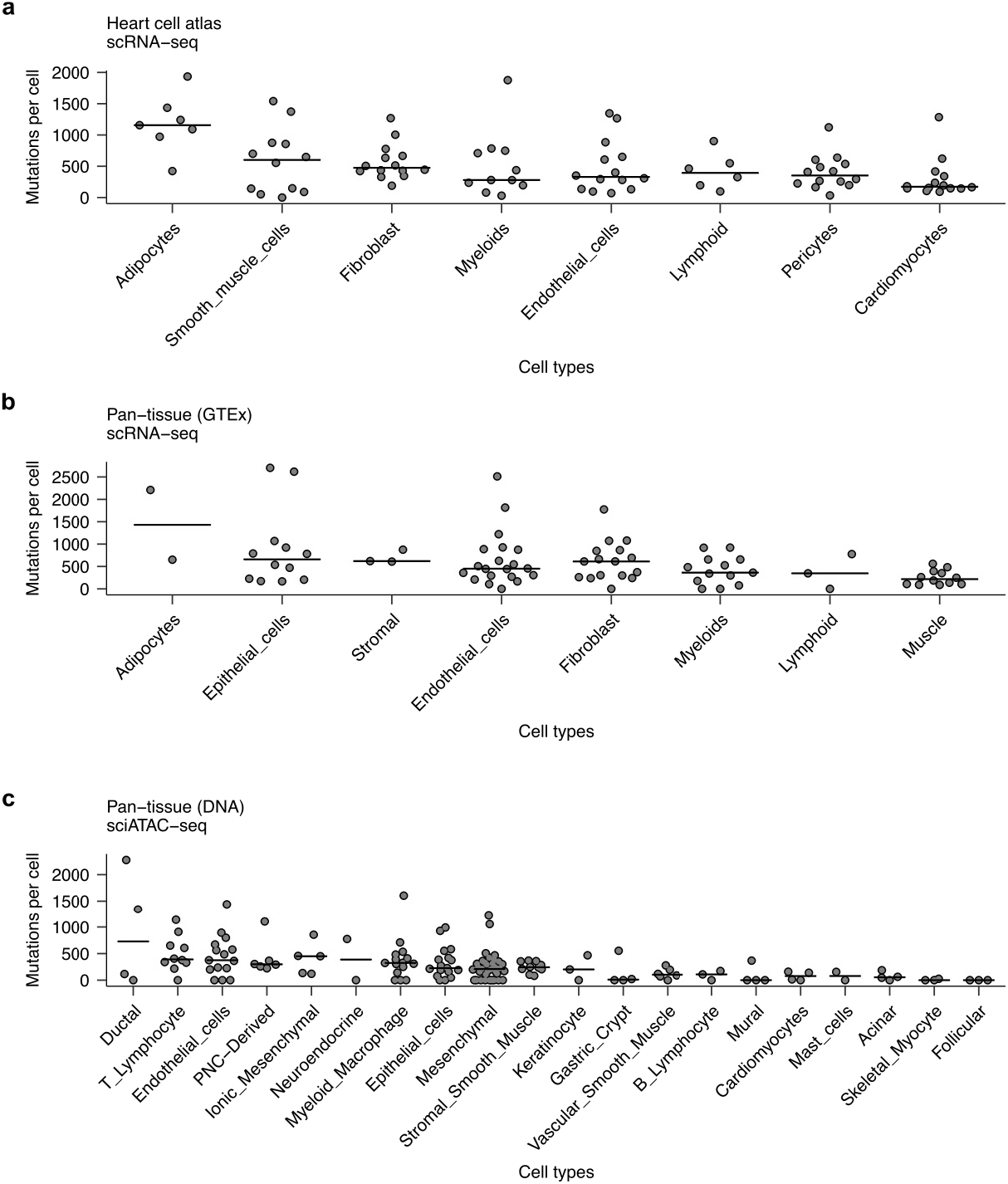
Mutational burdens estimated for single cells. Average mutational burden for single cells across the cell types detected in the scRNA-seq data from **(a)** the heart cell atlas, **(b)** pan-tissue GTEx, and **(c)** pan-tissue sciATAC-seq data sets. Each dot represents the average number of mutations estimated for each cell per sample. Only samples with at least 100 cells per cell type and datasets with at least two samples are shown. The horizontal line shows the median value across samples.

**Supplementary Figure 9.**
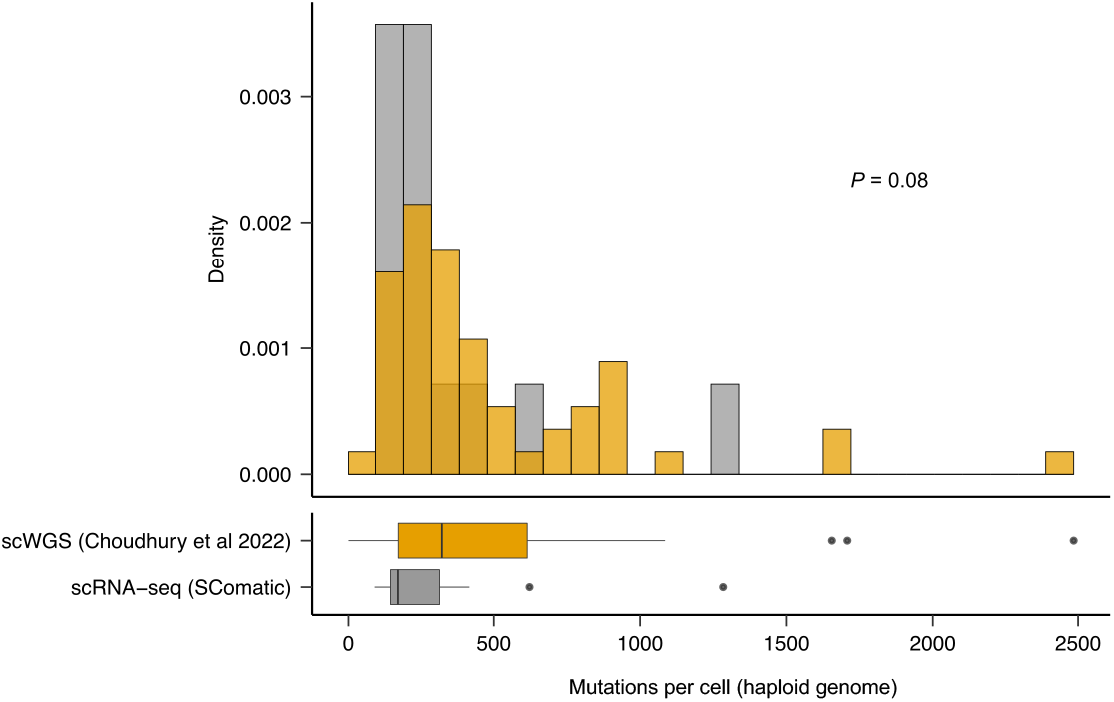
Rate of SNVs in cardiomyocytes computed using scRNA-seq from the Heart Cell Atlas and scWGS data from Choudhury *et al*. (*Nature Aging*, 2022). Mutation burdens were normalised to mutations per Mb and are expressed as mutations per cell and haploid genome. The mutation burdens estimated using scWGS data by Choudhury *et al*. (*Nature Aging*, 2022) were divided by the ploidy of each cell. The *P* value was computed using the two-sided Wilcoxon’s test.

**Supplementary Figure 10.**
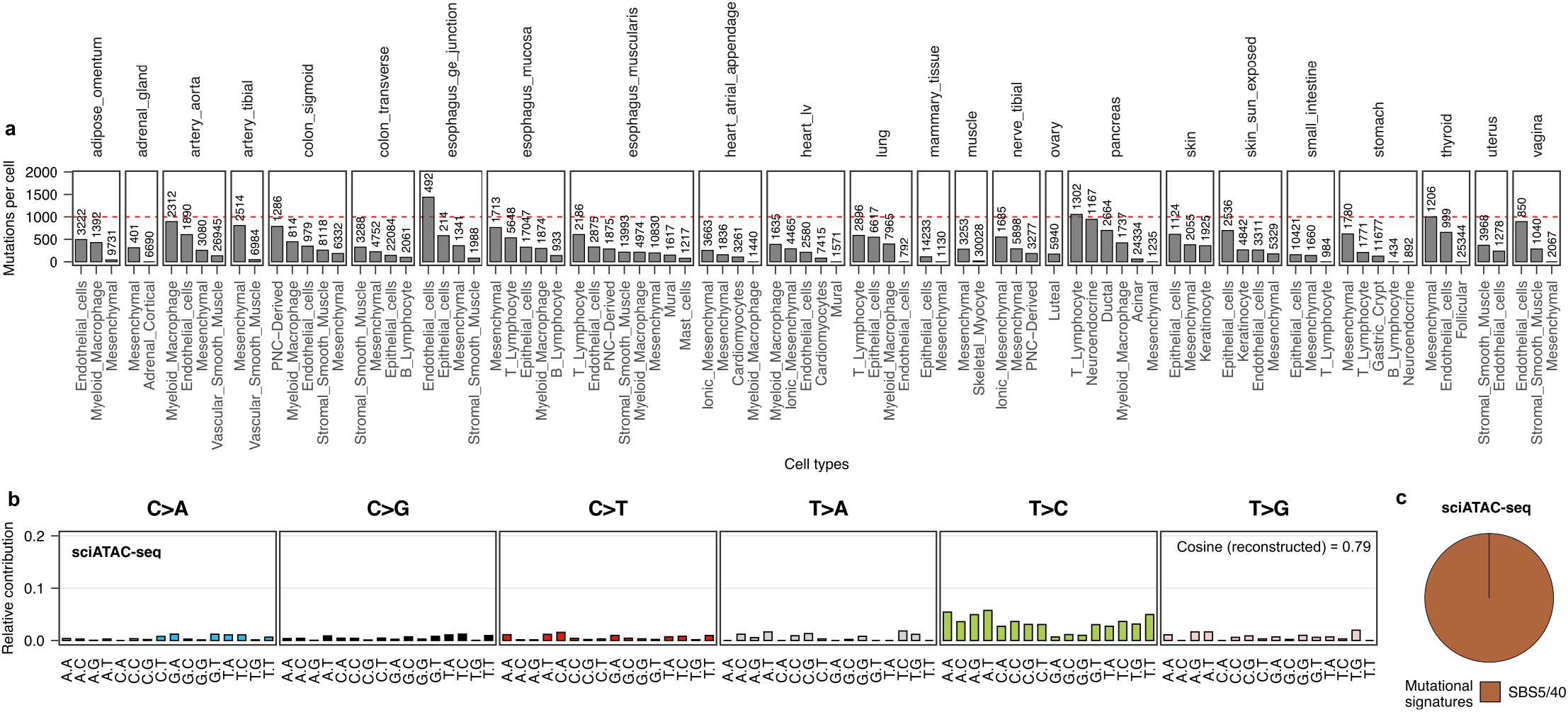
Somatic mutation detection in sciATAC-seq data. **a**) Average mutational burden at the single cell level estimated using the somatic mutations detected by SComatic in sciATAC-seq data. The mutational burden is expressed as mutations per cell and haploid genome. The number on top of the bars indicates the number of cells per cell type. **b**) Trinucleotide context of mutations detected across all cell types in the sciATAC-seq dataset. **c**) Decomposition of the mutations detected in sciATAC-seq data across all cell types into COSMIC signatures (reconstructed cosine similarity = 0.79). The contributions of SBS5 and SBS40 are collapsed for visualization purposes.

**Supplementary Figure 11.**
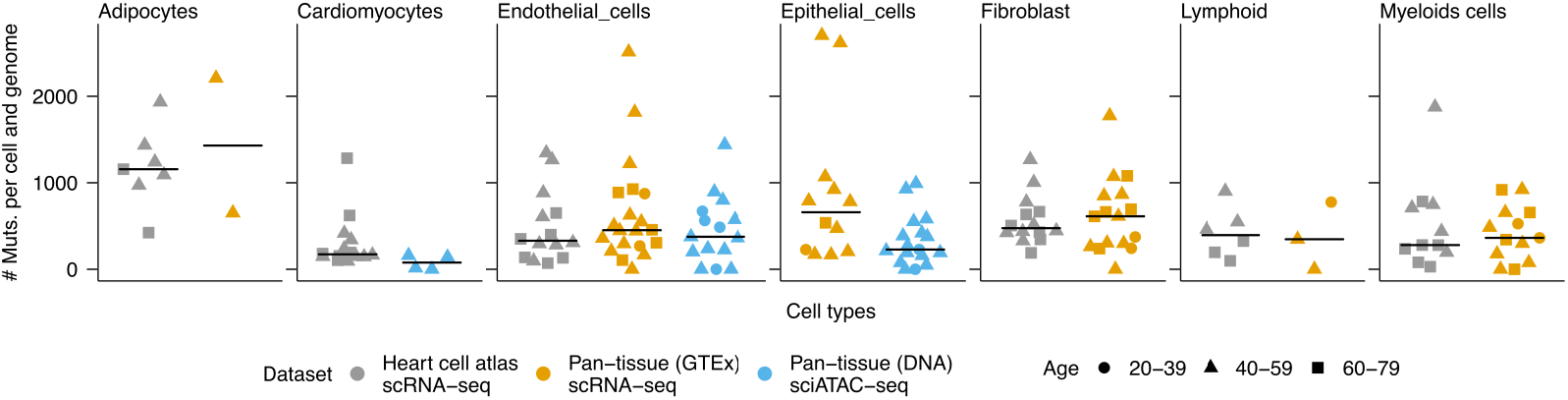
Comparison of the mutational burdens estimated for single cells across datasets. Each dot represents the average number of mutations detected per cell and haploid genome for each donor. The horizontal line shows the median value across samples. Only datasets with at least two samples and cell types present in at least two datasets are shown.

**Supplementary Figure 12.**
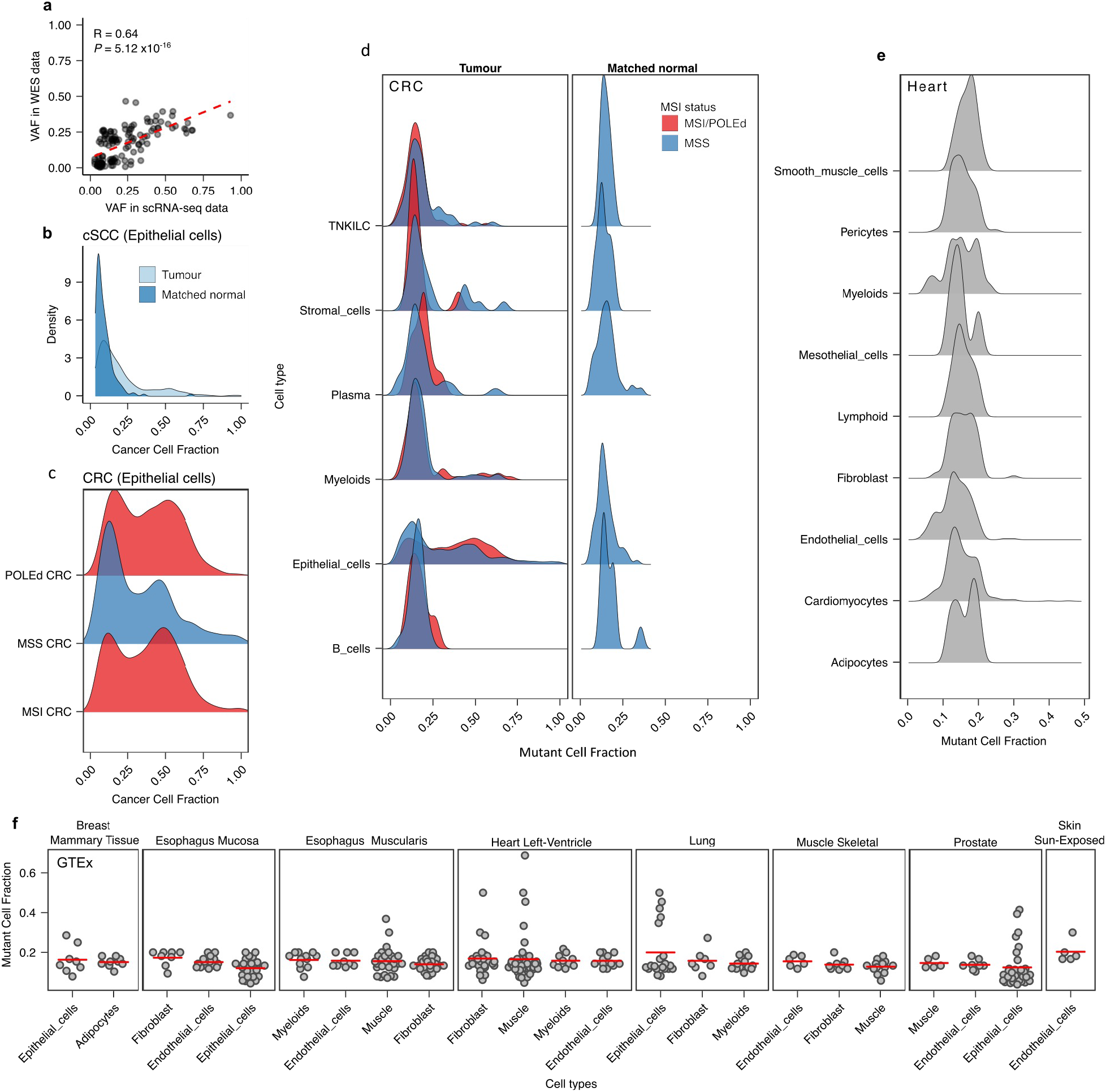
Clonality of the mutations detected in scRNA-seq data. **a**) Pearson correlation between the VAF of somatic mutations in WES and scRNA-seq data from the cSCC samples. Distribution of the cell fraction of mutations detected in scRNA-seq data from **b**) cSCC tumours and matched normal skin samples, **c**) colorectal tumours and matched normal samples, and **d**) epithelial cells from colorectal tumour samples. Distribution of the cell fraction of mutations detected across cell types from **e**) the heart cell atlas, and **f**) the GTEx data set. Each dot in **f** represents an individual SNV and the red horizontal line shows the mean for each group.

**Supplementary Figure 13.**
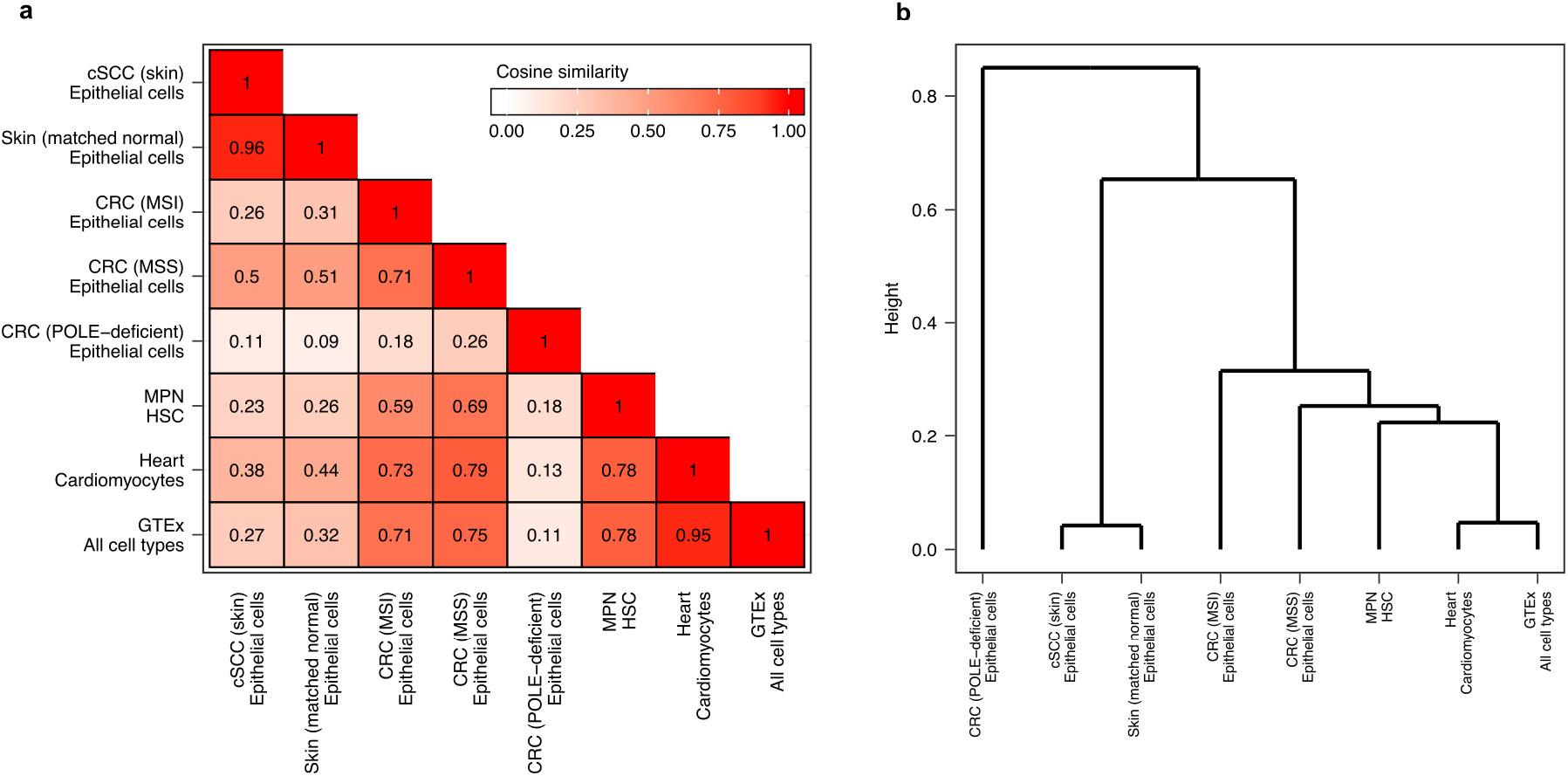
Comparison of the mutational patterns detected across the data sets analysed in this study. **a**) Pairwise cosine similarities between the mutational spectra computed using the mutations detected across cell types from each data set. **b**) Hierarchical clustering based on the cosine similarity comparison (shown in **a**) of the mutational spectra detected in each data set using SComatic.

**Supplementary Figure 14.**
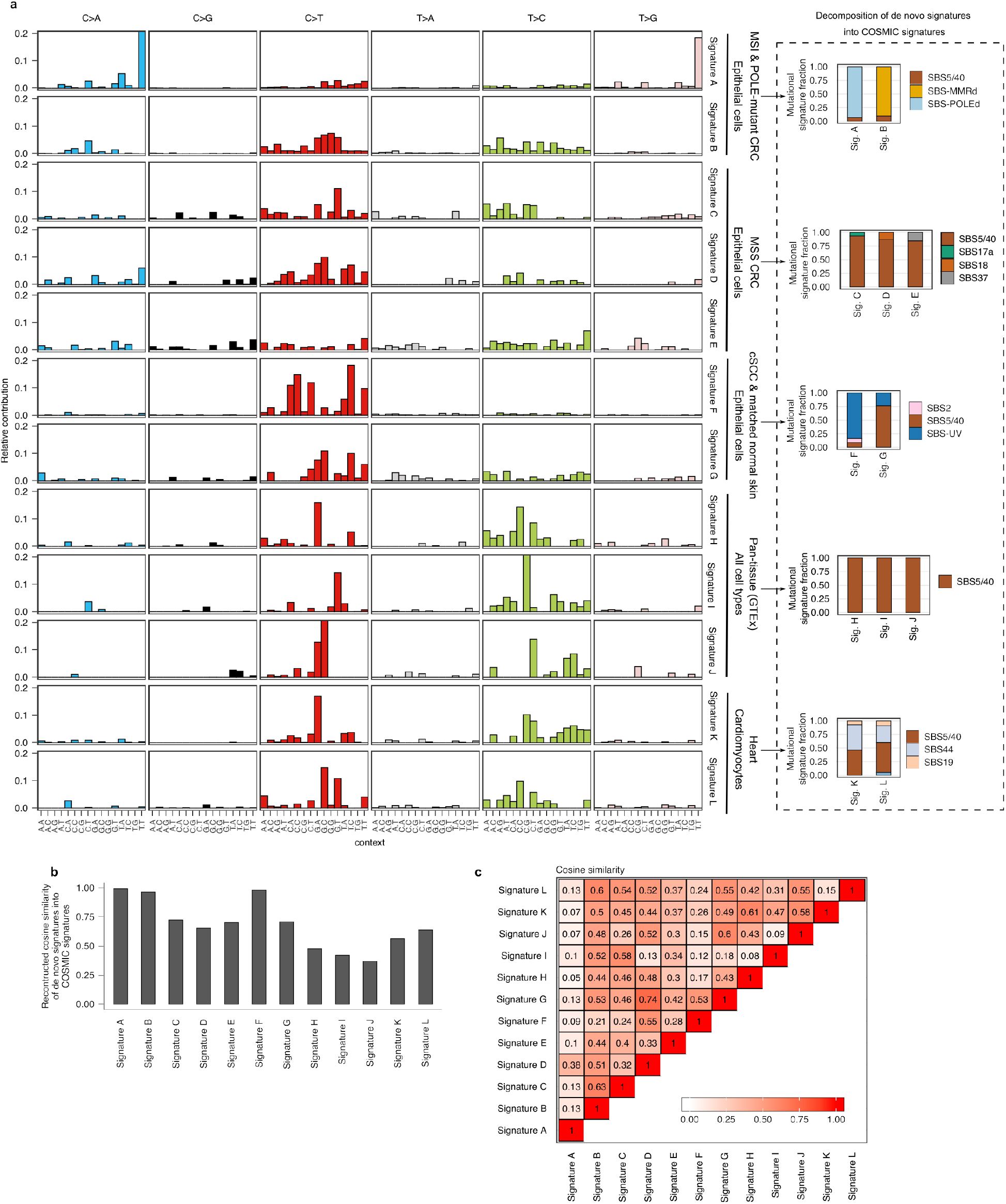
*De novo* mutational signature analysis of the somatic mutations discovered by SComatic. **a**) Trinucleotide context of the *de novo* signatures discovered in the data sets analysed. *De novo* mutational signatures were extracted independently from each dataset. The decomposition of the *de novo* signatures into COSMIC signatures was also run for each dataset independently. **b)** Cosine similarities between the *de novo* signatures and the reconstructed mutational spectra using the estimated signature contributions. **c**) Pairwise cosine similarities between each pair of mutational signatures extracted *de nov*o. Mutational signatures associated with MMRd (SBS6, SBS14, SBS15, SBS21, SBS26 and SBS44), POLE deficiency (SBS10a, SBS10b and SBS28), ultraviolet radiation (SBS7a,b,c and d) and clock-like mutational processes (SBS5 and SBS40) are collapsed for visualization purposes.

**Supplementary Figure 15.**
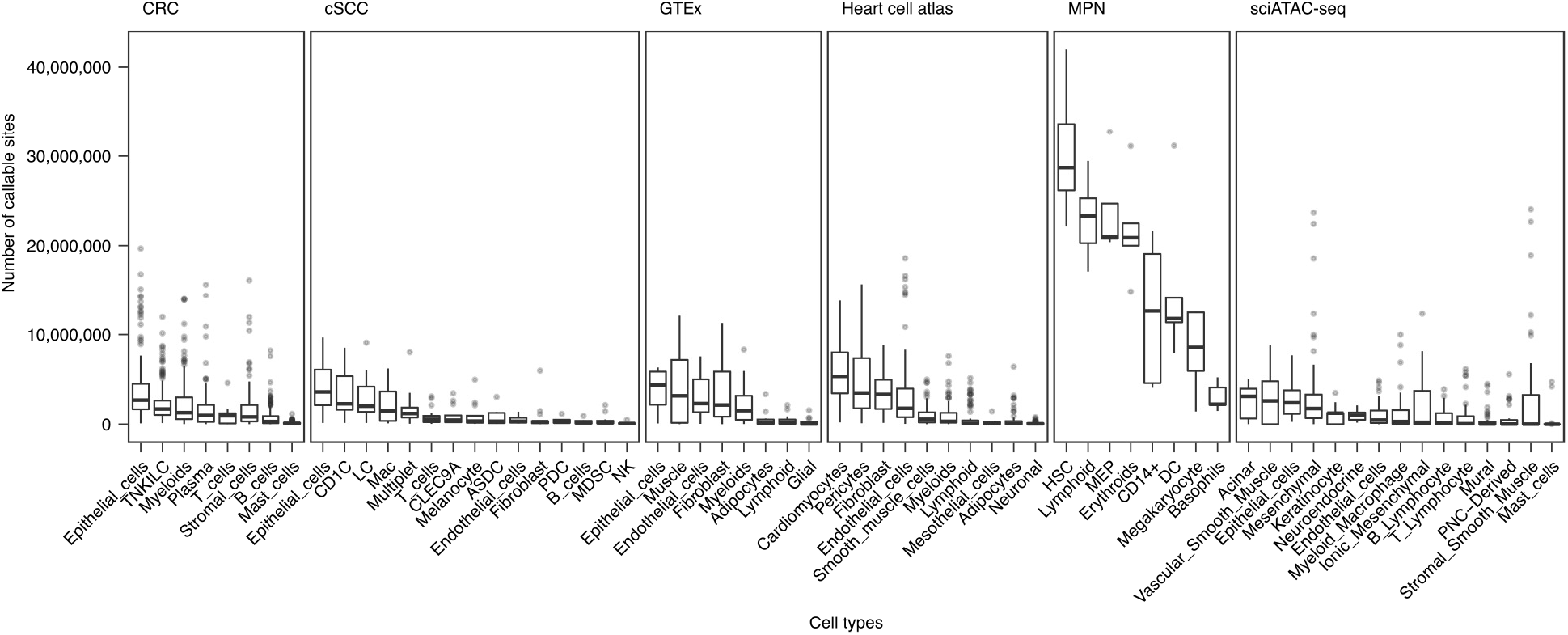
Number of callable sites per cell type and data set.

## Supplementary Tables

**Supplementary Table 1.**
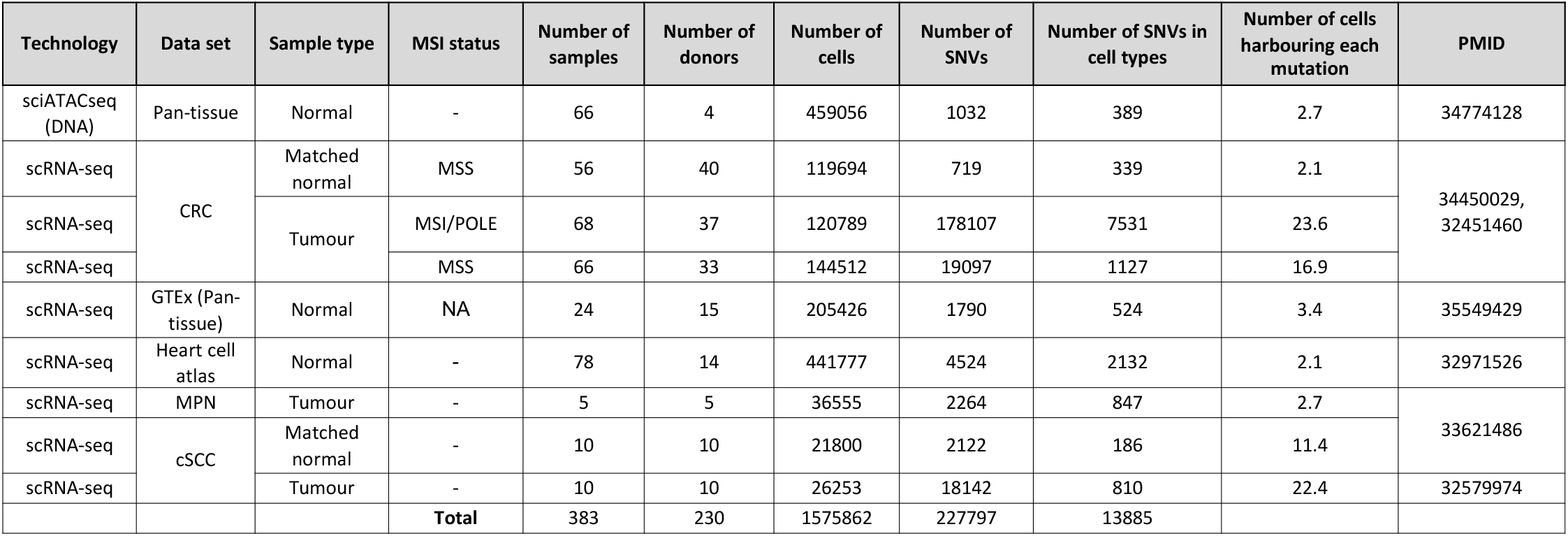
Summary of the data analysed and the somatic mutations detected in each data set.

